# Aardvark: Sifting through differences in a mound of variants

**DOI:** 10.1101/2025.10.03.680257

**Authors:** James M. Holt, Zev Kronenberg, Christopher T. Saunders, Egor Dolzhenko, Peter Krusche, Nathan D. Olson, Justin M. Zook, Michael A. Eberle

## Abstract

Variant benchmarking is critical in assessing the accuracy of genomic secondary pipelines. However, traditional benchmarking tools that require exact genotype matches inject biases from variant representation and are ill-suited for tandem repeat or structural variation. We describe Aardvark, a variant benchmarking tool that introduces the basepair score to directly compare haplotype sequences, reducing representation biases while allowing for partial credit scoring. The tool also includes a traditional genotype score and supports separate or joint benchmarking of small variants, tandem repeats, and structural variants (<10 kb). Aardvark accepts standard inputs, runs ≈16x faster than hap.py, and is freely available and open source (https://github.com/PacificBiosciences/aardvark).

## 1 Background

Variant benchmarking is paramount in assessing, validating, and improving sequencing technologies and variant calling pipelines. For clinical settings, measuring the performance of variant calling pipelines through benchmarking is necessary for establishing the accuracy and limitations of the technology and methodology. Genomic secondary pipelines often conclude by calling variants from reads aligned to a reference genome, which are typically reported in one or more Variant Call Format (VCF) files. The generated VCF files, or “query” sets, are then compared against well-established “truth” or “benchmark” sets to evaluate the accuracy of the final product. The earliest genome-scale benchmarks from Genome in a Bottle (GIAB) focused on non-repetitive genomic regions without structural variation, where variants are relatively easy to consistently detect across a wide range of technologies [22]. However, technical limitations restricted these early benchmarks to primarily unphased small variants, limiting the scope of benchmarking that could be performed. Recent benchmarks have been extended to generate sequence-resolved haplotypes that span more difficult regions of the genome [20, 15, 11] by leveraging a mixture of complementary sequencing technologies, advancements in variant calling and *de novo* assembly algorithms, and pedigree inheritance information. For example, the GIAB Telomere-to-Telomere (GIAB-T2T) [15] and Platinum Pedigree (PlatPed) [11] benchmarks include phased small variants, tandem repeats, and structural variants (≥ 50 bp). For variant benchmarking, these sequence-resolved haplotypes are reduced to variants relative to a reference genome, and the benchmarks are provided as a set of one or more VCF files with associated high-confidence regions. When applying these benchmarks, the Global Alliance for Genomic Health (GA4GH) benchmarking standards recommend stratifying the results by variant type, variant size, and by the genomic context around the variant to best characterize the performance and limitations of a query set [12].

Small variant benchmarking is still the most common form of assessment, specifically focusing on single-nucleotide variants (SNVs) and small insertions or deletions (“indels”, typically <50 basepairs). These small variants are detectable by many types of genomic assays, have been extensively studied by the community, and many have well-established phenotypic outcomes. Arguably, the *de facto* standard tool for performing small variant benchmarking is hap.py [12] with vcfeval [2] as the comparison engine performing the underlying assessments. The hap.py pre-processing step normalizes variants, which helps reduce the comparison errors due to differences in how a variant (or group of variants) are represented between the truth and query callsets. Vcfeval uses a dynamic program to maximize the number of detected true positives in the dataset, and includes logic such that the variant representations in each set do not have to precisely match so long as a matching pair of identical haplotype sequences generated by the variant calls can be found. Ultimately, any matching variants between the truth and query sets are labeled as true positives (TP), and the mismatches are labeled as false negatives (FN) or false positives (FP), respectively. Hap.py uses a genotype-centric approach for calculating summary metrics, meaning that labels are assigned one per genotype in the VCF, and those labels are used to calculate recall, precision, and F1 scores.

However, there are limitations and biases that are implicit to genotype-centric approaches. First, all variants are scored equally regardless of the size of the change. For example, a 1-bp indel and a 50-bp indel have the same representative weight despite the latter impacting 50x more sequence and thus having an increased likelihood to be deleterious in an individual [7]. This genotype-centric approach may best represent accuracy in coding regions, where it is often critical for variant and genotype calls to be exactly correct. For other applications and genomic contexts (e.g., larger variants outside of coding regions or in repetitive regions), it may be better to instead compare variants by their difference in sequence, affording larger events an increased relative weight since they impact more of the genome. Second, equivalent representations can be scored differently despite encoding the same underlying genetic changes. For example, some genomic contexts allow for a 2-bp insertion to be represented as a single 2-bp entry in a VCF or two nearby 1-bp entries. With genotype-centric scoring, the former will only be scored once whereas the latter representation is scored twice, implicitly doubling the weight of the latter representation. This can create biases either for or against some variant representations, leading to different outcomes and summary metrics even when the variants represent sequence-identical haplotypes [4].

Traditional genotype-centric scoring also does not allow for partial credit, which is becoming increasingly relevant as the benchmarks shift towards sequence-resolved haplotypes in repetitive regions and with larger forms of variation. Inexact allele matches often occur around larger indels, where precisely identifying the full variant sequence has more potential for error, particularly in homopolymers and tandem repeats. Outside of small variant benchmarking, structural variant callers frequently produce variant calls that are slightly different from a benchmark call, and the benchmarks themselves may include inexact endpoints for the structural variants. Similarly, tandem repeat variants have historically been analyzed by just looking at the expansion or contraction length, but recent studies have indicated sequence composition is also relevant for tandem repeat analysis [19]. Tools like Truvari [8] use partial credit to measure the similarity of these larger forms of variation, and typically use a heuristic cut-off (e.g., 70% sequence similarity) to determine if two variants count as true positive, false positive, or false negative. Vcfdist [4, 5] assigns partial credit based on the sequence similarity of the alleles, supporting both small variants and medium-sized structural variants (<10 kb). For example, a 9-bp homopolymer insertion compared to a 10-bp insertion would receive a score of 0.9, as opposed to 0.0 from a traditional pass/fail score. While both Truvari and vcfdist use partial credit to assist in matching and scoring variants, they are both still genotype-centric approaches, assigning equal weight to each genotype in the file and leaving them subject to similar representation biases as hap.py/vcfeval. Vcfdist additionally requires that both the truth and query variants are locally phased to perform the comparison. These challenges have pushed some researchers to shift from variant-based evaluations to sequence-based evaluations. In particular, the recent Telomere-to-Telomere “genome benchmark” and accompanying software (“Genome Quality Checker”) compare genome-scale sequences (i.e., assemblies) instead of variants [9]. With a sequence-based approach, the summary metrics are instead based on the number of true or false basepairs detected in a query sequence, enabling implicit partial credit relative to a genotype-based approach. While genome benchmarking can assess accuracy of genomes inferred from fully-phased variant calls, the process of converting variant calls to genomes is challenging, and there is still a need for sequence-centric metrics that assess variant accuracy directly.

We introduce “Aardvark”, a benchmarking tool that implements a sequence-centric scoring scheme suitable for small variants, tandem repeats, and medium-sized structural variants (<10 kb). This “basepair” scoring scheme reconstructs the benchmark haplotype sequences, generates query haplotype sequences, and then compares them at the basepair level. The basepair scheme is implicitly variant-type agnostic, weights each modified basepair equally, and significantly reduces representation biases that are inherent in a genotype-centric approach. Aardvark also includes a traditional genotype scoring scheme that mimics hap.py, assigning equal weight to each genotype in the truth and query sets regardless of the zygosity or length of the variant. The tool ingests standard file formats and generates summary metrics, labeled VCF files, per-region metrics, and stratification results for each run. Aardvark is also computationally efficient, generating all of the above outputs ≈16x faster than hap.py, enabling faster iteration when testing new variant calling methods.

## 2 Results

### 2.1 Aardvark for variant benchmarking

Aardvark is a benchmarking tool that efficiently evaluates the accuracy of a query set against a benchmark set using both a sequence-centric approach and a traditional genotype-centric approach. The tool groups variants by proximity into a set of sub-regions that are evaluated independently and in parallel. In each sub-region, the tool calculates an optimal phase orientation of the provided variants, generates localized haplotype sequences, and evaluates the result with the sequence-centric (“basepair” or `BASEPAIR`) scoring scheme and the genotype-centric (“genotype” or `GT`) scoring scheme (see Figure 1). Aardvark is open source and publicly available, and we have released several examples showing similarities and differences with genotype-based benchmarking tools (see Supplemental Materials). We demonstrate the versatility of Aardvark by testing multiple combinations of publicly available truth benchmarks [22, 20, 15, 11] in conjunction with query datasets [1, 18] generated from a diverse set of sequencing technologies and variant calling pipelines (see Supplemental Materials).

**Figure 1:**
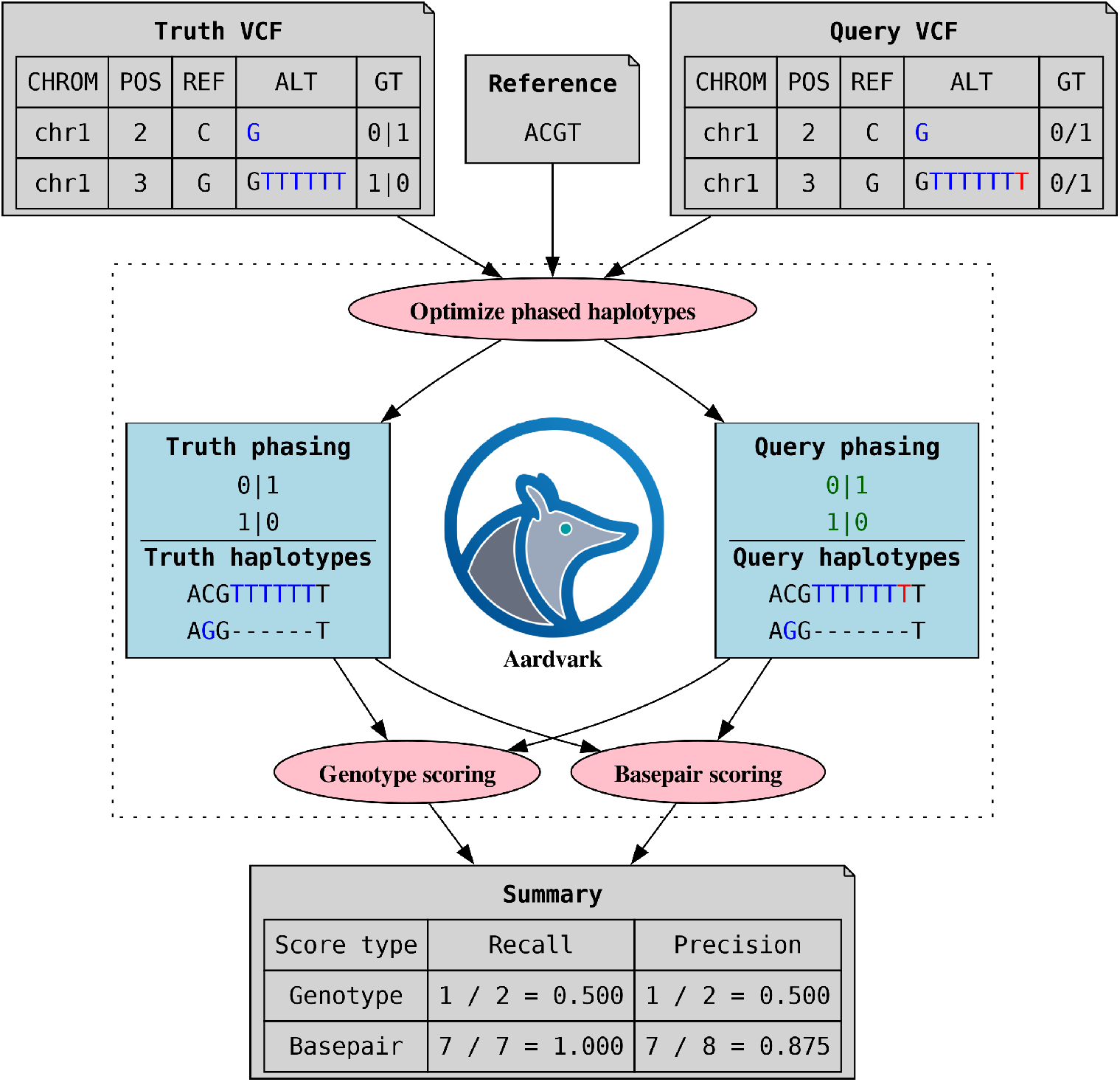
Aardvark process overview. Aardvark ingests a reference genome file (FASTA) and two VCF files: one for truth variants and one for query variants. It then calculates the optimal phasing of the query variants such that the edit distance between the truth and query haplotypes is minimized. Once the optimal phasing is identified, the phased variants are scored using a genotype scoring scheme. Independently, the haplotype sequences are also scored with the variant-agnostic basepair scoring scheme. Metrics from both scores are provided in a summary file. In this example, the truth and query VCFs share a single-nucleotide variant, but the insertion variant in the query VCF (7T) is one basepair longer (red) than the one in the truth VCF (6T). For genotype scoring, this one basepair difference creates both a false positive and false negative for the insertions, leading to recall and precision of 0.500. With basepair scoring, the only difference is a single false positive “T” base in the first haplotype, leading to a recall of 1.000 with a precision of 0.875.

### 2.2 Aardvark generates sequence-centric summary metrics

Aardvark’s basepair scoring scheme shifts from a genotype-centric approach to a sequence-centric approach. With basepair scoring, the variants are masked by inserting them into localized haplotype sequences, and then comparing those sequences at the basepair level. Since the variants are masked, the summary metrics are instead calculated by identifying the number of true positive and false positive/negative basepairs between the truth and query haplotypes. Thus, all summary metrics for the basepair scoring scale with the number of altered basepairs within the VCF files instead of the number of genotype calls. Larger variants implicitly have weight proportional to their length, which may give the appearance of up-weighting these variants relative to a genotype-centric approach. However, Aardvark’s basepair scoring actually equalizes the weight of each altered basepair, and biases from variant representation are significantly reduced if not removed entirely (see Methods).

Compared to Aardvark’s genotype scores, the overall basepair scores are lower for short-read sequencing query sets (mean F1 delta: − 0.0070) but marginally higher for long-read sequencing query sets (mean F1 delta: +0.0004). This pattern is primarily driven by indel scoring, where F1 scores decrease for short-read query sets (mean delta: − 0.0169) but increase for long-read query sets (mean delta: +0.0183, see Figure 2). When we inspected the detailed results, we found that the short-read query sets excel at shorter indels (≤ 15 bp) but struggle with longer indels (>15 bp), an observation that is supported by both hap.py and Aardvark-`GT` labeled outputs (see Supplemental Materials). These longer indels alter more basepairs and therefore have increased relative weight with basepair scoring. This culminates in basepair scores that are lower than the genotype scores for these short-read query sets. In contrast, the long-read query sets had better performance with the longer indels, leading to increased F1 scores even when the reported alleles included minor sequence errors. For long-reads, these relevant increases were most prominent in tandem repeat and homopolymer regions (see Supplemental Materials). We note that these trends persist with short-read and long-read query sets from the Precision FDA Truth Challenge v2 [16] (see Supplemental Materials).

**Figure 2:**
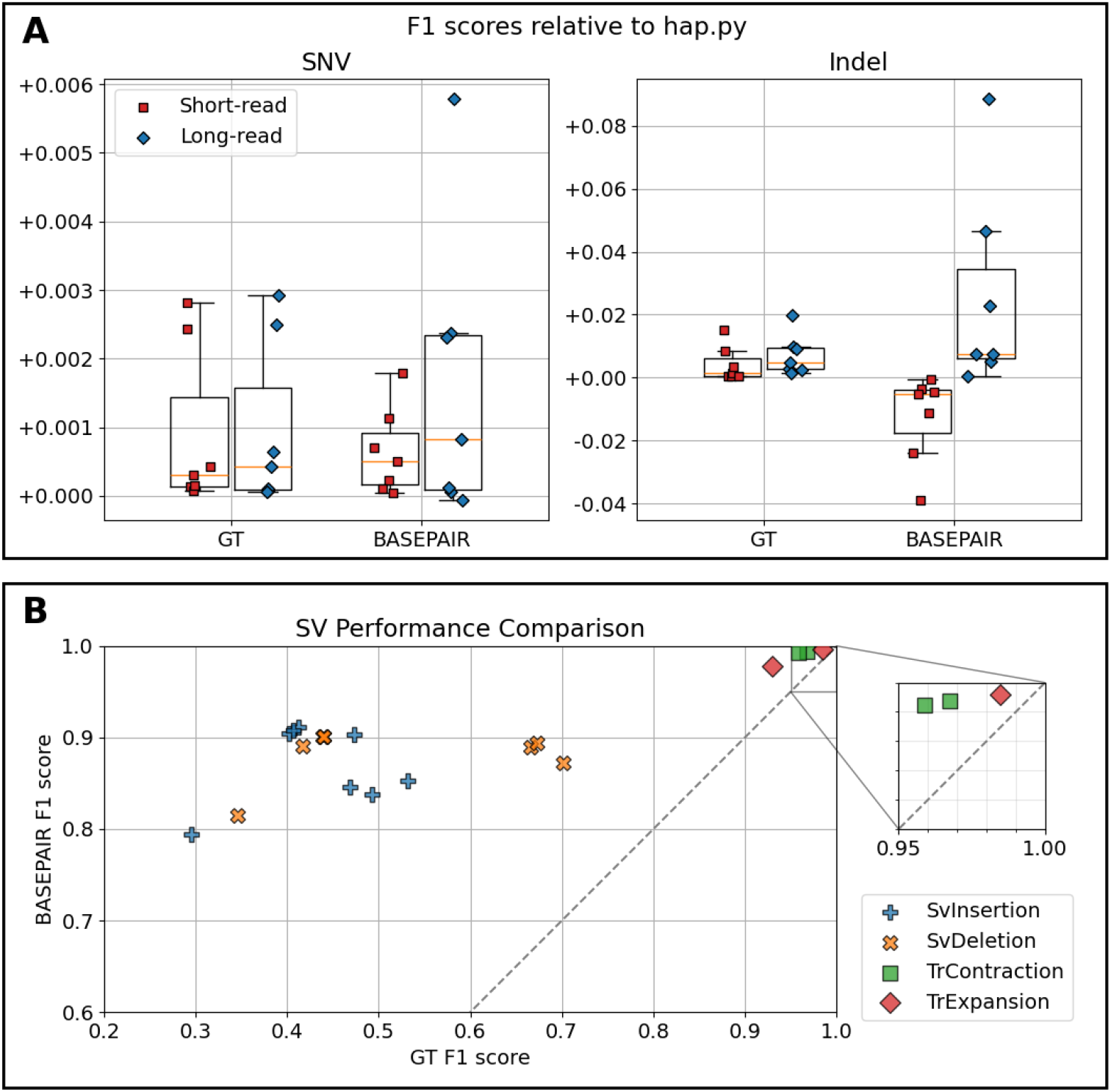
**(A)** F1 scores of small variants relative to hap.py. These figures show the difference in F1 scores relative to hap.py for SNVs (left) and indels (right) for Aardvark’s genotype (`GT`) and basepair (`BASEPAIR`) scoring schemes. The truth sets include GIAB-v4 (HG001/HG002), PlatPed (HG001), GIAB-CMRG (HG002), and GIAB-T2T (HG002). The query datasets are divided into short-read and long-read sequencing categories. On average, the genotype and basepair scores increase slightly for SNVs regardless of sequencing type. For the short-read query sets, the indel basepair scores decrease due to difficulties calling longer indels (>15 bp). In contrast, the basepair scores for long-read query sets improve due to increased ability to capture the longer indels. **(B)** F1 scores of tandem repeat and structural variants in Aardvark. This table shows the genotype and basepair F1 scores for various long-read tandem repeat and structural variant calls against either the PlatPed (HG001) or GIAB-T2T (HG002) truth sets. The variant callers identify the majority of modified bases, which is reflected in the relatively high basepair F1 scores. However, the reported variant alleles often do not exactly match the truth set alleles, leading to much lower genotype F1 scores for the same comparison.

### 2.3 Aardvark benchmarks tandem repeat and structural variants

Aardvark’s basepair scoring is well-suited for benchmarking medium-sized (<10 kb) tandem repeat or structural variant calls, which are much more likely to include minor sequence errors due to their length. To demonstrate this capability, we compared multiple tandem repeat or structural variant query sets from long-read sequencing against the Platinum Pedigree [11] and GIAB Telomere-to-Telomere [15] benchmarks, restricting both the variants and regions to match the query sets. Tandem repeats showed a modest improvement when using Aardvark’s basepair F1 scores instead of genotype F1 scores (mean delta:+0.0297, see Figure 2). For structural variants, the basepair F1 scores were substantially higher than the genotype F1 scores (mean delta: +0.4011), highlighting how structural variant callers frequently find the majority of modified bases but do not always generate an exact allelic match to a truth set (see Figure 2). This effect is most pronounced with structural variant insertions (mean delta:+0.4344) where the variant callers must assemble the inserted sequence.

### 2.4 Basepair scoring reduces representation biases

Aardvark’s basepair scoring compares sequences and is inherently variant-agnostic, significantly reducing biases from alternate variant representations. To demonstrate this capability, we leveraged benchmark sets from the Platinum Pedigree [11] which include separate small variant and tandem repeat call sets. While the variants and confidence regions for these benchmark sets are very different, we identified ≈ 413k tandem repeat regions for which the variants generate identical haplotype sequences in each benchmark set, making them theoretically interchangeable. For query sets, we selected two long-read sequencing datasets with matching runs of both a small variant caller (DeepVariant [17]) and a tandem repeat caller (TRGT [3]), and then compared each pair of truth and query set using Aardvark.

For a given query set, we observe that the basepair F1 scores are always identical, even though the truth sets have different variant representations (see Table 1). In contrast, the genotype scores change when the truth sets use different representations, especially if the query set consists of small variants. Additionally, the basepair F1 scores are consistently higher than the corresponding genotype F1 scores, highlighting how sequence-level comparisons with partial credit often boosts F1 scores for larger forms of variation. This increase is most prominent with the tandem repeat truth and query sets (+0.0444 and +0.0164 relative to genotype F1), where any sequence-level mismatch will often lead to multiple false labels with genotype scoring.

**Table 1:**
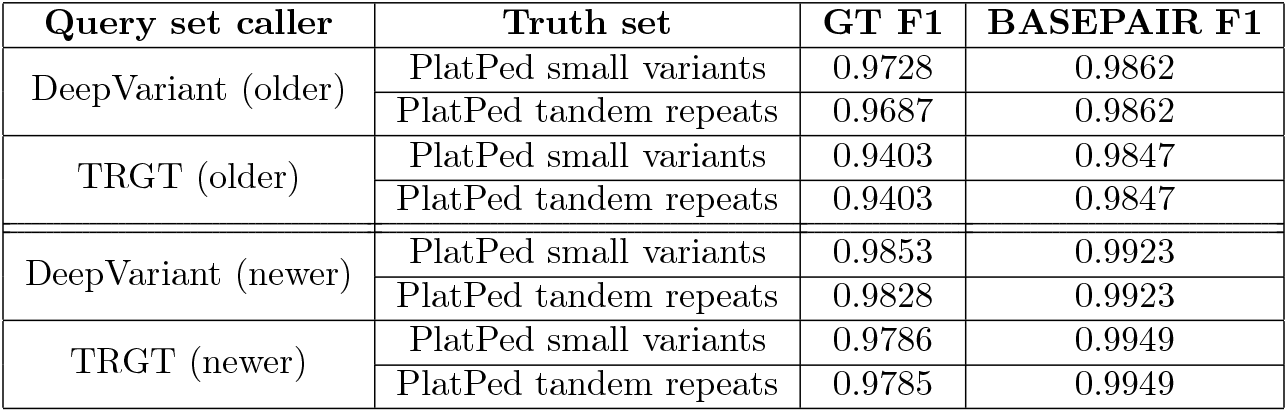
Aardvark comparisons in Platinum Pedigree tandem repeat regions. This table shows the F1 score from Aardvark for different combinations of truth and query sets restricted to tandem repeat regions. For query sets, we used DeepVariant (small variant caller) and TRGT (tandem repeat caller) calls from the same HG001 long-read datasets. We selected an older dataset with matching older runs for each of the tools, and a newer dataset with newer runs of each tool. For truth sets, we used the small variant and tandem repeat benchmarks from the Platinum Pedigree (HG001), limiting the analysis to regions that are sequence-identical in each set (≈ 413k repeat regions). The genotype F1 scores (“GT F1”) show variability in the results due to representation biases and binary pass/fail scoring. In contrast, the basepair F1 scores are identical for a query set regardless of the comparator truth set, and they are consistently higher than the genotype F1 scores due to sequence-level comparisons with partial credit for inexact matches.

### 2.5 Aardvark for joint benchmarking of different variant classes

Aardvark can be used to jointly benchmark multiple classes of variation at the same time if fully integrated VCFs are provided for the truth and query sets. We demonstrate this with the full GIAB Telomere-to-Telomere benchmark [15] which includes both small and structural variants as a joint benchmark set. For a query set comparator, we ran an experimental joint variant calling pipeline that produces integrated small and structural variants in a single VCF file (see Supplemental Materials). We compared these integrated variant sets using Aardvark, first limiting the comparison to the small variant benchmark regions and then with the complete region set. While the structural variants make up only 1.32% of the joint benchmark genotypes, they cover 65.06% of the altered basepairs in the benchmark, which translates into a roughly 3-fold increase in the total number of basepairs relative to the small variants alone.

With the joint comparison, Aardvark calculates a structural variant genotype F1 score of 0.8740 and a basepair F1 score of 0.9130, leading to overall drops in the F1 scores when considering all variants (see Table 2). The impact on the overall genotype F1 score is low (−0.0048) since there are relatively few structural variants in the dataset (1.32%). For basepair scoring, the structural variants carry significantly more weight (65.06%), leading to a larger drop in the overall F1 score (−0.0438) when all variants are considered together. Additionally, the computational cost of the increased scope of comparisons (3-fold increase) is reflected in the increased compute time for Aardvark (from ≈1 minute to ≈3 minutes, see Table 2).

**Table 2:** Example joint Aardvark comparisons. This table shows multiple Aardvark comparisons of the GIAB-T2T benchmark against a query set generated from an experimental joint variant calling pipeline based on *de novo* assembly. Both the truth and query sets include integrated small and structural variants in a single VCF file. The first comparison is limited to the GIAB-T2T small variant regions, whereas the second and third comparisons are against the full GIAB-T2T benchmark which includes more challenging regions with structural variants. For both GIAB-T2T region sets, we show the percentage of benchmark variation that is structural (“Percent SV”) from both genotype (`GT`) and basepair (`BP`) perspectives. The first two comparisons used Aardvark’s default window size of 50 bp, and we include a third comparison that increases it to 1 kb. The F1 scores from both genotype and basepair scoring is shown for all variants and structural variants alone (“SVs F1”). The last column shows Aardvark’s wall clock time to complete each comparison with 16 threads.

| T2T regions | Percent SV |  | Window size | All variants F1 |  | SVs F1 |  | Wall clock time (m:ss) |
| --- | --- | --- | --- | --- | --- | --- | --- | --- |
|  | GT | BP |  | GT | BP | GT | BP |  |
| Small variants | 0.00% | 0.00% | 50 | 0.9842 | 0.9874 | – | – | 0:51.26 |
| Full benchmark | 1.32% | 65.06% | 50 | 0.9794 | 0.9436 | 0.8740 | 0.9130 | 3:19.15 |
|  |  |  | 1000 | 0.9802 | 0.9546 | 0.8964 | 0.9181 | 9:35.75 |

We found that increasing Aardvark’s window size parameter led to improved accuracy scores for the full benchmark comparison. In Aardvark, increasing the window size creates larger sub-problems, which can have theoretical benefits for matching equivalent variants with distant representations. By increasing the parameter from the default of 50 bp up to 1 kb, we increase the overall basepair F1 score by +0.0110 but at the cost of increased compute time (from ≈3 minutes to ≈10 minutes, see Table 2). We do not observe this F1 score improvement with only small variant comparisons (see Supplemental Materials), suggesting that the difference is caused by the inclusion of structural variants and/or the more challenging regions containing them.

### 2.6 Aardvark accelerates variant benchmarking

Aardvark computes and reports all results very efficiently using less than 3 minutes of wall-clock time for each small variant comparison in our analyses. Aardvark used 6.0% of the total wall clock time of hap.py, making it ≈16x faster (see Figure 3). Within this limited compute time, Aardvark produces basepair scoring metrics and traditional genotype scoring metrics that are highly concordant with hap.py (99.63-99.99%, see Supplemental Materials). Aardvark also allows for stratification of the results based on user-provided regions, enabling more focused accuracy metrics but with increased compute costs. The differences in compute requirements are more pronounced when we enable stratifications, with one test case requiring over an hour for hap.py while the equivalent Aardvark run finished in ≈2 minutes. Aardvark’s max memory usage across all non-stratified tests was ≈18 GB, increasing to ≈26 GB for the stratified tests. While our method of running hap.py could not precisely measure the memory requirements, we had to allocate over 32 GB to complete the non-stratified runs which increased to over 64 GB when stratifications were enabled. Overall, these metrics indicate that Aardvark generates all benchmark results and output files efficiently, using a fraction of the compute resources of hap.py.

**Figure 3:**
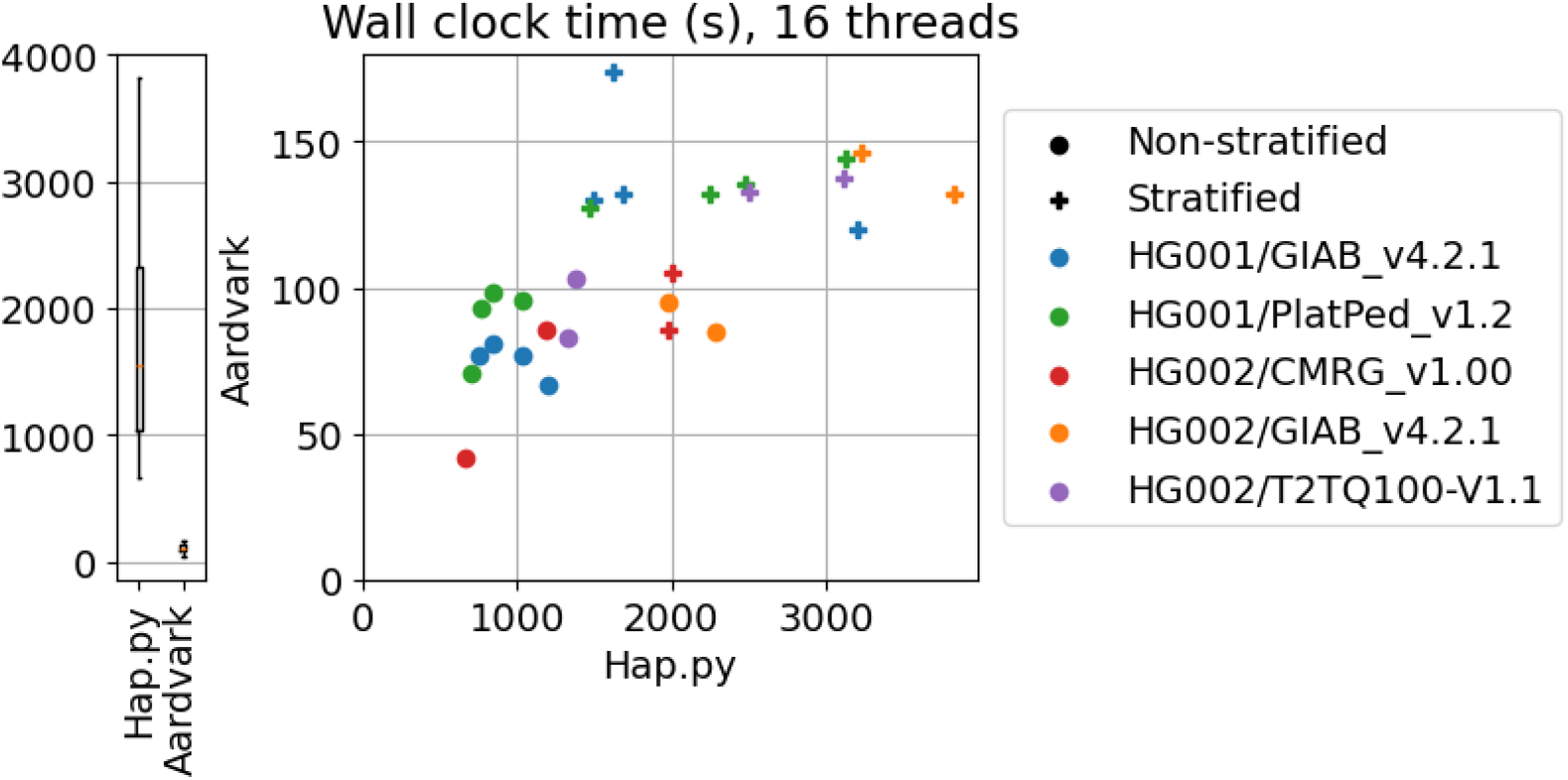
Comparison of wall-clock compute times with 16 threads. Left: The distribution of wall-clock compute times for hap.py and Aardvark. Right: Scatter plot of the same data demonstrating the impact of the benchmark type and stratification on the run times. Enabling stratifications increases the runtime of all comparisons (189 stratifications covering 219 million total regions). In total, Aardvark used 6.0% of the wall-clock time of hap.py.

## 3 Discussion

Aardvark’s basepair score provides a sequence-level comparison that address many of the critiques of genotype scoring. Specifically, basepair scoring 1) reduces or removes biases from variant representation which may be caused by technology or tooling, 2) equalizes the weight of each modified basepair, 3) affords additional weight to homozygous calls and allows for partial credit for zygosity errors, and 4) enables partial credit when allelic sequences do not exactly match (see Supplemental Materials for detailed list). Basepair scoring is analogous to the “genome benchmark” approach [9], but it operates on smaller local haplotypes derived from VCF files and does not require fully-phased variant calls for the benchmark and query sets. The genotype score is intended to serve as a replacement for hap.py with practical enhancements in the form of improved speed and greater ability to resolve complex variation. We consider these two scoring schemes to be complementary, with basepair scoring providing an approach that is much closer to complete haplotype-resolved benchmarking, whereas genotype scoring may be more suited for applications where strict exact-matching variants are critically important such as evaluation of variant calls in clinically-relevant regions.

Aardvark’s basepair scoring scheme provides a benchmarking view that may be useful for improving variant callers. First, sequence-level comparisons enable benchmarking approaches that would be less accurate with a genotype score. For example, we demonstrated how Platinum Pedigree tandem repeats and small variants benchmarks can be used interchangeably to achieve the same basepair score metrics, enabling developers to perform benchmarks that were previously inaccessible due to significant differences in variant representation. Second, the increased resolution from basepair scoring enables stratification of regions by their haplotype correctness. This can help developers categorize errors by type, sequence composition, and/or severity, enabling more focused assessment and development of tools. Lastly, variant callers that leverage machine learning [17, 21, 1] could be modified to use the basepair scoring metrics as part of the optimization error to minimize. A genotype-based score may get caught in local minima where improvements to imperfect allelic sequences may manifest as identical scores. Basepair scoring could allow the training to escape these local minima through the detection of partially correct results, which may lead to faster convergence or increased accuracy in the final variant calling models.

We demonstrated Aardvark’s ability to jointly benchmark different classes of variation, but there are not many available resources or variant callers that can fully leverage this capability. To our knowledge, the GIAB-T2T benchmark is the only available truth set with integrated small and structural variant calls in a single file. All other benchmarks in our experiments split the variants into different files, allowing for focused comparisons of a class of variation but at the cost of potentially duplicate or conflicting variants. Similarly, variant callers (i.e., query sets) are often specialized to a class of variants, which can lead to conflicting and overlapping variant calls that require additional computation to resolve. Recent assembly-based variant callers will report all variant types in one phased VCF without conflicts [13, 6], but these callers have additional limitations from assembly. While comprehensive joint benchmark resources and tools may not be quite ready for routine use, we hope to leverage the capabilities of Aardvark to benchmark their outputs and identify areas for future improvement.

## 4 Conclusion

Aardvark is a new variant benchmarking tool that introduces the basepair scoring scheme, enabling direct haplotype sequence comparisons and reducing biases from variant representation. Additionally, Aardvark supports joint benchmarking of small variants, tandem repeats, and structural variants, enabling more comprehensive comparisons with assembly-based benchmarks and variant callers. The implementation is computationally efficient while reporting both the basepair score and a traditional genotype score. The tool ingests standard file formats and outputs labeled variants, summary statistics, and several optional files for users who desire deeper details on each individual haplotype comparison. Aardvark is open source, easy to install, and freely available on GitHub (https://github.com/PacificBiosciences/aardvark).

## 5 Methods

### 5.1 Overview

Aardvark prioritizes a sequence-centric approach to variant benchmarking. Thus, the core algorithmic problem is to efficiently compare diploid haplotype sequences that are derived from two sets of input variants: “truth” and “query” sets. To take advantage of parallelism, Aardvark first divides the problem space into sub-regions that are solved independently. For each sub-region, Aardvark will 1) construct optimized truth and query haplotype sequence pairs from the provided variants such that the total edit distance between the pairs is minimized, and 2) score the optimization solution using multiple scoring schemes.

### 5.2 Region identification

Performing a full end-to-end comparison for an entire chromosome is computationally challenging. Additionally, truth sets often come with “confidence” or “benchmark” regions indicating where the comparison should be performed for the highest accuracies, effectively excluding regions where the truth sets do not contain high-confidence variation. Aardvark starts with these confidence regions, and further divides them into sub-regions based on the input variants and a user-defined window heuristic (default: *W* = 50 bp). When parsing the truth and query variants, any variants from either input that are within *W±* basepairs of each other are grouped together into a sub-problem. These sub-problems are analyzed independently, allowing for Aardvark to leverage parallelization by distributing the sub-problems to different threads. This windowing scheme is similar to approaches taken by other tools to sub-divide the problem [12, 4]. Increasing the window size creates larger sub-problems and may improve accuracy under certain conditions such as when larger variants are part of the benchmark, but it also increases the computational complexity of each sub-problem. When restricted to only small variants, we found negligible differences in the summary F1 scores beyond the default 50 bp window size, and the additional compute requirements started to become noticeable beyond 200 bp (see Supplemental Materials). For structural and tandem repeat variants, we found 1 kb to be a suitable window size for most applications. While this heuristic window size approach is computationally fast, more sophisticated strategies like those of vcfdist [5] may prove to be better for accuracy.

### 5.3 Sequence optimization

For each sub-problem, Aardvark starts by constructing two pairs of truth and query haplotype sequences (four total) that minimize the total edit distance between each truth/query sequence pair. These sequences are constructed using the variants and zygosities provided in the truth and query variant files. A homozygous variant must be incorporated into both corresponding haplotype sequences (e.g., a truth variant must go into both `truth1` and `truth2`). However, a heterozygous variant is incorporated into exactly one of the two haplotypes, and the phase orientation defines which haplotype incorporates the variant (e.g., a truth variant with orientation 1|0 will be incorporated into `truth1`, but not `truth2`). Thus, the core problem is to find an optimal combination of *phased* zygosities such that the generated haplotype sequences have the smallest combined edit distance (e.g., minimize `ed(truth1, query1) + ed(truth2, query2)`).

When phased genotypes are provided for the truth variants, Aardvark will respect them, meaning that the algorithm will not alter or “flip” the phase orientation of any phased truth zygosities. If the truth variants are unphased, the algorithm is allowed to flip their phase orientations to minimize the edit distance calculations. In contrast, query phase orientations are never respected as it is assumed that phasing errors are possible in the provided data. For most modern truth sets, the variants are all phased onto maternal and paternal haplotypes, so this routine is typically only searching for the optimal query phase orientations.

Given *V* total unphased heterozygous variants, there are 2^*V*^ possible phase orientations that exist. Aardvark models this search space as a decision tree where at depth *V* the phase of the *V* th variant (ordered by position) is assigned. Aardvark explores this decision tree using a lowest-cost-first algorithm based on the total edit distance between the truth and query haplotype sequences represented by the node. The method is initialized at the root node (*V* = 0) with no assigned zygosities. Each possible phase orientation is explored and added to the minimum cost queue using the edit distances derived from comparing the partial haplotypes. This process is repeated until depth *V* is reached, indicating that all variants have been phased and a minimum-cost solution has been found. Conceptually, this process is similar to the phasing algorithm described for HiPhase [10], but with a slightly altered search space and optimization scheme adapted for this problem.

The underlying edit distance calculations are handled by a custom dynamic version of the wavefront algorithm [14] (WFA). The base WFA algorithm is very efficient, with run-time scaling based on the edit distance between two sequences. Aardvark’s dynamic version allows for the pair of sequences (truth and query) to each be extended as the search algorithm traverses down the decision tree. Additionally, this method does not apply a penalty if there are extra characters in the partially constructed sequences, as future extensions may lead to matches. Given that the vast majority of haplotypes are expected to exactly match for high-quality variant calling pipelines, this method of exploring the search space often traverses directly down the search tree to the correct answer in *O*(*V*) time, with negligible computation cost from the WFA calculations. However, regions with higher error rates can lead to much more branching (*O*(2^*V*^)) with higher WFA costs, thus Aardvark bounds the search by limiting the number of nodes searched at each depth to a fixed heuristic value (*H*), limiting the worst case search to *O*(*HV*). In practice, we found that Aardvark rarely hits these limits, but when it does it is usually in areas that are enriched for false positives and/or negatives (i.e., the truth/query haplotypes have significant divergence).

### 5.4 Scoring

After identifying the optimal phase orientation for all truth and query variants, Aardvark scores each pair of haplotypes independently (truth v. query) and combines the scores for the region. Aardvark scores the differences at the sequence (or haplotype) level, fully masking variant representation from consideration for reporting the results in the basepair (`BASEPAIR`) scoring scheme. The tool also reports genotype (`GT`) level metrics which are highly similar in both method and outcome to those reported by previous methods [2, 12]. Aardvark also reports two other genotype-based scores with alternate weights (see Supplemental Materials).

#### 5.4.1 Basepair scoring scheme

Aardvark’s basepair scoring scheme directly compares two haplotypes sequences without knowledge of the underlying variants. This approach starts by constructing haplotype sequences from the optimized truth and query variant phasing, effectively blinding all downstream computations to the original variant representations. For both pairs of truth/query haplotypes, it counts the number of bases in the haplotype sequences that are true positives (changes shared in both truth and query relative to the reference sequence), false negatives (changes unique to truth), or false positives (changes unique to query). Thus, the denominator in metrics like precision and recall scaled with the total number of modified, inserted, or deleted basepairs relative to the reference sequence.

By definition, all differences between the reference genome sequence and a truth haplotype sequence must be either a true positive or a false negative. Similarly, each difference between the reference sequence and a query haplotype sequence must be either a true positive or a false positive. Lastly, each difference between the truth and query haplotypes must be either a false negative or false positive. Given a method to calculate the edit distance between two sequences, these definitions form a system of equations that can be trivially solved for each pair of haplotype sequences. Let *R* be the reference genome sequence in the region, *T* be the constructed truth haplotype sequence, and *Q* be the constructed query haplotype sequence. Let the edit distances be calculated by some function, *ed*(*A, B*), between each pair of input sequences: *X* = *ed*(*R, T*), *Y* = *ed*(*R, Q*), and *Z* = *ed*(*T, Q*). Finally, let *TP* be the number of true positives, let *FN* be the number of false negatives, and let *FP* be the number of false positives. Given these definitions, we create the following system of equations where *TP, FN*, and *FP* are initially unknown, and all the edit distances are calculated from the optimized haplotype sequences:

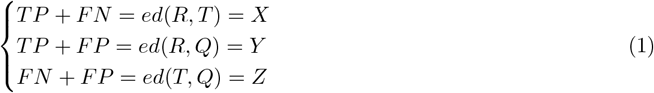

Which simplifies to:

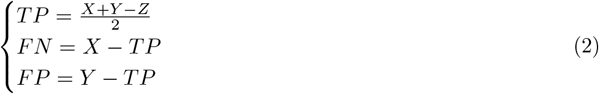

This system of equations is solved twice, once for each pair of optimized truth and query haplotype sequences. The edit distance calculations are the most costly part of this process, which is implemented using the wavefront algorithm [14] to minimize run-time. In practice, this approach is highly efficient since the vast majority of comparisons are exact matches. Interestingly, these definitions also allow for TP, FN, and FP values that are half-correct (i.e., include a 0.5 fractional component). These half-correct bases typically represent situations where the truth and query both incorporate alternate alleles, but with slightly different sequences. In practice, Aardvark doubles all basepair scoring counts (TP, FN, and FP) to remove floating-point calculations as a source of technical error. Table 3 includes several examples demonstrating the doubled base counts and partially correct sequences.

**Table 3:** Examples of differences between Aardvark’s genotype (`GT`) and basepair (`BASEPAIR`) scoring schemes. The examples in this table include the reference sequence (REF), the truth sequence, the query sequence, and the generated number of true positives (TP), false negatives (FN), and false positives (FP) for both genotype and basepair scoring schemes. To avoid floating-point numbers from “half-correct” base changes, `BASEPAIR` values are twice the number of altered basepairs. Set A are all heterozygous variants with a single base change, including two simple examples where the “half-correct” changes are relevant in basepair scoring (odd numbers are caused by “half-correct” bases). Set B includes at least one homozygous variant in each set, highlighting how zygosity errors are scored with each approach. Set C are all heterozygous insertions, showing major scoring differences when partial credit is applied in homopolymer indel regions. Set D includes a mix of heterozygous and homozygous genotype calls on alleles that reflect common errors in homopolymer variant calling. FN and FP are always unique to truth and query, respectively. For genotype scoring, TP may be different between truth and query (impacting recall and precision) and is annotated as “Truth.TP” and “Query.TP” where relevant. If TP, FN, or FP is unlisted, it is =0.

| Set | REF | Truth | Query | GT | BASEPAIR |
| --- | --- | --- | --- | --- | --- |
| A | A | A/C | A/C | TP=1 | TP=2 |
|  | A | A/C | A/A | FN=1 | FN=2 |
|  | A | A/A | A/C | FP=1 | FP=2 |
|  | A | A/C | A/G | FN=1, FP=1 | TP=1, FN=1, FP=1 |
|  | A | A/C | A/AC | FN=1, FP=1 | TP=1, FN=1, FP=1 |
| B | A | C/C | C/C | TP=1 | TP=4 |
|  | A | C/C | A/C | FN=1, Query.TP=1 | TP=2, FN=2 |
|  | A | C/C | A/A | FN=1 | FN=4 |
|  | A | A/C | C/C | Truth.TP=1, FP=1 | TP=2, FP=2 |
|  | A | A/A | C/C | FP=1 | FP=4 |
| C | A | A/ACCC | A/ACCC | TP=1 | TP=6 |
|  | A | A/ACCC | A/ACC | FN=1, FP=1 | TP=4, FN=2 |
|  | A | A/ACCC | A/ACCCC | FN=1, FP=1 | TP=6, FP=2 |
| D | A | C/C | C/AC | FN=1, Query.TP=1, FP=1 | TP=3, FN=1, FP=1 |
|  | A | ACCC/ACCC | ACC/ACCCC | FN=1, FP=1 | TP=10, FN=2, FP=2 |

The core benefit of the basepair scoring scheme is that variant representation biases have been removed entirely. First, the total weight of any comparison is always equal to the number of altered basepairs, removing biases from zygosity or variant representation. Second, equivalent variant representations have identical summary recall values when errors are present (see Table 4). This enables greater resolution in the summary metrics while also significantly reducing biases that may be injected from alternate variant representations.

**Table 4:** Examples of representation biases. This table shows multiple simplified examples of variant combinations that are labeled with a count, zygosity (heterozygous or homozygous), and variant type. The variant combinations are grouped into sets based on the number of altered basepairs (changed, inserted, or deleted), highlighting situations that could be sequence-identical. For both the genotype (`GT`) and basepair (`BASEPAIR`) scoring schemes, the total weight of the variants under that scoring scheme is shown along with the recall metric if a 1-bp false negative error was present in the query set. Set A has a basic heterozygous SNV where the weights and scores are identical. Set B shows two scenarios that equal 2 basepairs of inserted sequence, emphasizing how zygosity can influence the weight in each score. Sets C and D show four scenarios each that equal a 50 bp heterozygous or homozygous indel, further emphasizing major differences in weighting and scoring between the approaches. The total weight of the genotype score is always equal to the number of variant entries from the VCF file, while the basepair score weights are equal to the number of modifed basepairs. The basepair weights and scores do not change within the equivalent sets, whereas genotype scoring is heavily influenced by the variant representation and zygosity.

| Set | Variants | Altered bp | GT | BASEPAIR |
| --- | --- | --- | --- | --- |
| A | One heterozygous SNV | 1 | 1 (0.0) | 1 (0.0) |
| B | Two heterozygous 1-bp indels | 2 | 2 (0.5) | 2 (0.5) |
|  | One homozygous 1-bp indel | 2 | 1 (0.0) | 2 (0.5) |
| C | One heterozygous 50-bp indel | 50 | 1 (0.0) | 50 (0.98) |
|  | Two heterozygous 25-bp indels | 50 | 2 (0.5) | 50 (0.98) |
|  | Ten heterozygous 5-bp indels | 50 | 10 (0.9) | 50 (0.98) |
|  | Fifty heterozygous 1-bp indels | 50 | 50 (0.98) | 50 (0.98) |
| D | One homozygous 50-bp indel | 100 | 1 (0.0) | 100 (0.99) |
|  | Two homozygous 25-bp indels | 100 | 2 (0.5) | 100 (0.99) |
|  | Ten homozygous 5-bp indels | 100 | 10 (0.9) | 100 (0.99) |
|  | Fifty homozygous 1-bp indels | 100 | 50 (0.98) | 100 (0.99) |

#### 5.4.2 Genotype scoring scheme

For the genotype scoring scheme, Aardvark starts with the same phase orientations from sequence optimization, but uses an orthogonal algorithm to compute the metrics. For each pair of alternate alleles, Aardvark searches for the maximum number of alternate alleles that can be incorporated into the reference sequence such that the output truth and query haplotype sequences exactly match. Critically, the allele representations do not have to match, so long as the final haplotype sequences are basepair-identical. Any alleles that are incorporated are marked as true positives, and all others are either false negatives or false positives depending on the source. For homozygous genotypes, the genotype score further requires that both alleles are marked as true positives for the entire genotype to also get marked as a true positive. In general, this method is consistent with previous approaches and the outcomes are highly concordant with hap.py (see Supplemental Materials).

There are two subtleties worth mentioning about the Aardvark implementation of the genotype score. First, any multi-allelic heterozygous sites are split into two distinct heterozygous genotypes for all analyses. Second, if the difference between truth and query is limited to the zygosity of the genotype call (i.e., heterozygous v. homozygous), then the heterozygous call is labeled as “true” and the homozygous call is labeled “false”. For example, if the truth set has a heterozygous call while the query set has homozygous, then the truth variant is labeled as a true positive since it was correctly detected in the query, but the query variant is labeled as a false positive since an extra allele is present. Both of these subtleties are distinctions from the hap.py genotype scoring [12], and are sources of technical discrepancy when comparing the two tools (see Supplemental Materials).

### 5.5 Truth and query datasets

Throughout this manuscript, we used a variety of publicly available benchmarks (truth sets), all of which are for the well-studied HG001 (NA12878) or HG002 samples. For HG001 benchmarks, we used Genome in a Bottle (GIAB) v4.2.1 [22] and Platinum Pedigree (PlatPed) v1.2 [11]. For HG002 benchmarks, we used Genome in a Bottle (GIAB) v4.2.1 [22], GIAB Challenging Medically Relevant Genes (GIAB-CMRG) v1.00 [20], and GIAB Telomere-to-Telomere (GIAB-T2T) Q100 v1.1 [15]. The GIAB v4.2.1 sets are relatively strict and focus mostly on non-repetitive genomic regions, meaning they are typically “easier” to resolve correctly with modern sequencing technologies and variant callers. In contrast, the CMRG, PlatPed, and GIAB-T2T benchmarks all include more difficult regions of the genome, which are typically much harder for modern pipelines to resolve correctly.

For query sets, we leveraged a combination of short-read and long-read datasets from previous publications [1, 18] and other public resources. These query sets are generated using a variety of pipelines and sequencing technologies. While efforts were made to select high quality variant calls for these comparisons, this is a rapidly developing field and the approaches used to generate these variant calls may be outdated for a variety of reasons. While the Aardvark tool is intended to analyze the accuracy of variant calling, the analyses provided in this document were chosen to demonstrate the capabilities of the tool and the effect of alternate scoring schemes in practice, and they should *not* be used to compare the underlying technologies or pipelines. As such, each query set is either completely anonymized or labeled with the corresponding read-length type (short-read or long-read), as this was the only relevant feature we identified for understanding the differences between Aardvark’s scoring schemes. Links to the exact files for our benchmark and de-anonymized query sets are available in our Supplemental Materials.

## 6 Acknowledgments

NDO and JMZ received support from NIST intramural funding. Certain commercial equipment, instruments, or materials are identified to specify adequately experimental conditions or reported results. Such identification does not imply recommendation or endorsement by the National Institute of Standards and Technology, nor does it imply that the equipment, instruments, or materials identified are necessarily the best available for the purpose.

## Notes

https://github.com/PacificBiosciences/aardvark

https://doi.org/10.5281/zenodo.17227384

https://github.com/holtjma/mini_variant_benchmarks

### Competing Interest Statement

JH, ZK, CS, ED, and ME are all current employees of PacBio. PK is a current employee of Novartis Pharma AG.

