## Supplemental Materials for "Aardvark: Sifting through differences in a mound of variants"

James M. Holt et al.

October 3, 2025

### 1 Data collections

#### 1.1 Data bundle

Some variant files were pre-processed to prevent crashes in hap.py and/or Aardvark. This typically removed homozygous reference calls and extraneous calls from outside the benchmark regions (i.e., ALT contigs or similar), filtered the variants to autosomes, and/or altered poorly formatted VCF lines to comply with VCF specification. To maximize reproducibility of the analyses we performed, the exact pre-processed truth and query VCF files, BED regions, GIAB stratifications, and both tools (hap.py and Aardvark) are bundled into a single Zenodo URL: `https://doi.org/10.5281/zenodo.17227384`.

#### 1.2 Truth set collections

Table 1 lists each benchmark (truth set) used in our analyses. Each benchmark has an accompanying high confidence region BED file, which indicates where the comparisons should be restricted to. Additionally, a set of standard stratifications were downloaded from Genome in a Bottle (GIAB)[1] for testing the affect of stratification on performance. While the majority of these truth set files were unaltered, a few were pre-processed prior to benchmarking. See Section 1.1 to access the exact files used in our analyses.

| Variants | Sample | Benchmark | Reference | URL |
|---|---|---|---|---|
| Small | HG001 | Genome in a Bottle (GIAB) v4.2.1 | [12] | 2 |
| | | Platinum Pedigree (PlatPed) v1.2 | [5] | 3 |
| | HG002 | Genome in a Bottle (GIAB) v4.2.1 | [12] | 4 |
| | | Challenging Medically Relevant Genes (CMRG) v1.00 | [11] | 5 |
| | | Telomere-to-Telomere (T2T-Q100) v1.1 | [7] | 6 |
| Tandem repeats | HG001 | Platinum Pedigree (PlatPed) v1.2 | [5] | 7 |
| Structural | HG001 | Platinum Pedigree (PlatPed) v1.2 | [5] | 8 |
| Joint | HG002 | Telomere-to-Telomere (T2T-Q100) v1.1 | [7] | 9 |

Table 1: List of truth sets. A manuscript reference and download URL for each file has been provided. Some files were altered to resolve errors in hap.py. Refer to Section 1.1 to download the full set of files used after alterations.

#### 1.3 Small variant query set collections

Table 2 lists each small variant query set VCF used in our analyses. In general, these files represent relatively recent datasets that are broadly available to the public. References are listed when a dataset is tied to a publication, and brief descriptions of the prep, sequence, and pipelines are provided where possible. We note that some of these files were pre-processed prior to benchmarking. These datasets are unmasked here purely for reproducibility, and the analyses in this supplement or the main document are not intended to be nor should they be interpreted as formal comparisons between the sequencing technologies or tools. See Section 1.1 to access the exact files used in our analyses.

[1] `https://ftp-trace.ncbi.nlm.nih.gov/ReferenceSamples/giab/release/genome-stratifications/v3.6/`

[5] `https://ftp-trace.ncbi.nlm.nih.gov/ReferenceSamples/giab/release/NA12878_HG001/NISTv4.2.1/GRCh38/HG001_GRCh38_1_22_v4.2.1_benchmark.vcf.gz`

[6] `http://platinum-pedigree-data.s3.amazonaws.com/truthset_v1.2/NA12878_hq_v1.2.smallvar.vcf.gz`

[7] `https://ftp-trace.ncbi.nlm.nih.gov/ReferenceSamples/giab/release/AshkenazimTrio/HG002_NA24385_son/NISTv4.2.1/GRCh38/HG002_GRCh38_1_22_v4.2.1_benchmark.vcf.gz`

[8] `https://ftp-trace.ncbi.nlm.nih.gov/ReferenceSamples/giab/release/AshkenazimTrio/HG002_NA24385_son/CMRG_v1.00/GRCh38/SmallVariant/HG002_GRCh38_CMRG_smallvar_v1.00.vcf.gz`

[9] `https://ftp-trace.ncbi.nlm.nih.gov/ReferenceSamples/giab/data/AshkenazimTrio/analysis/NIST_HG002_DraftBenchmark_defrabbV0.019-20241113/GRCh38_HG2-T2TQ100-V1.1_smvar.vcf.gz`

| Sample | Sequencing technology | Reference | Description and URL |
|---|---|---|---|
| HG001 | Element Biosciences (SR) | [9] | Elevate library prep kit, AVITI 2×150bp flow cell, DeepVariant v1.6.1 variant caller; derived from 2188-E[10] |
| HG001 | Illumina (SR) | [1] | Illumina NovaSeq 6000 2×151 bp paired-end, DRAGEN 4.2.4[11] |
| HG001 | Oxford Nanopore Technology (LR) | – | ONT Kit V14, PromethION sequencer, Clair3 v1.0.0 variant caller[12] |
| HG001 | PacBio (LR) | [9] | SMRTbell prep kit 3.0, mixed Sequel IIe and Revio sequencers, DeepVariant v1.8.0 variant caller; derived from [13] |
| HG002 | Illumina (SR) | [1] | Illumina NovaSeq 6000 2×151 bp paired-end, DRAGEN 4.2.4; same URL as HG001 Illumina |
| HG002 | PacBio (LR) | – | SMRTbell prep kit 3.0, Revio sequencer, DeepVariant variant caller; re-analysis of [14] |

Table 2: List of small variant query sets. Where possible, we provide references, URLs to download the files, and details on the prep, sequencer, and pipelines used to generate each VCF. Some files were altered to resolve errors in hap.py. Refer to Section 1.1 to download the exact files used after alterations.

#### 1.4 Structural, tandem repeat, and joint query set collections

For structural, tandem repeat, and joint variant query sets, we limited the analysis to tools that could run on long-read sequencing datasets. Long reads are capable of generating sequence-resolved alleles through most medium-sized variation, and there are several tools available to generate these calls for structural or tandem repeat variation. We used HiFi sequencing technical replicates of HG001 and HG002, some from Sequel IIe and some from Revio. Each dataset was aligned with a standardized WDL workflow, but with possibly different versions of the underlying alignment or variant calling tools. For the variant callers, we ran pbsv[15], sawfish [10], TRGT [4], and/or longcallD[16] depending on the truth set variant compositions. See the data bundle for the exact files used in our analyses (Section 1.1).

#### 1.5 Benchmarking tool versions

Table 3 lists each benchmarking tool used in our analysis and the version. Additionally, a link to our miniaturized examples are included in the tools.

#### 1.6 Tool commands

All tools were run using a snakemake pipeline developed to automate the process.

##### 1.6.1 Hap.py

There are multiple ways to install and run hap.py in a cluster environment. Ultimately, we settled on using a Singularity (Docker) container that contained all the necessary components. We also tried running hap.py using a bioconda environment, but found that approach used significantly more memory than the Dockerized

[10] http://platinum-pedigree-data.s3.amazonaws.com/data/element/mapped/GRCh38/2188-E.GRCh38.merged.sort.bam
[11] https://zenodo.org/records/8350256
[12] http://ont-open-data.s3.amazonaws.com/giab_2023.05/analysis/variant_calling/hg001_sup_all/hg001.wf_snp.vcf.gz
[13] http://platinum-pedigree-data.s3.amazonaws.com/data/hifi/mapped/GRCh38/NA12878.GRCh38.haplotagged.bam
[14] https://downloads.pacbcloud.com/public/revio/2022Q4/HG002-rep1/
[15] https://github.com/PacificBiosciences/pbsv
[16] https://github.com/yangao07/longcallD

| Tool | Version | Description |
|---|---|---|
| Hap.py [6] | v0.3.8 | The main comparator tool in our analyses, and arguably the community standard for small variant benchmarking. This version is the Docker version tagged with "latest", which we have packed in a singularity image for reproducibility. GitHub: `https://github.com/Illumina/hap.py` |
| Aardvark | v0.7.4 | The novel tool we describe in the main document. All tests were performed using the pre-compiled static binary file that is distributed with each release. GitHub: `https://github.com/PacificBiosciences/aardvark` |
| Examples pipeline | – | We developed a pipeline to assist with both example generation and inspection of discordant results. The repo includes the scripts used to generate all output files, environments for running tools, the pipeline itself, and the outputs we have generated. See Section 2.3 for more details. GitHub: `https://github.com/holtjma/mini_variant_benchmarks` |

Table 3: Benchmarking tool versions and descriptions.

container (the reason for this is unknown). The downside of the Docker approach is that CPU time and memory consumption could not be accurately assessed using the snakemake benchmark functionality. The following command contains the exact parameterized options used to run hap.py:

```
/opt/hap.py/bin/hap.py \
    --threads {threads} \
    -o {params.out_prefix} \
    -r {input.reference} \
    -f {input.call_regions} \
    --gender {params.sex} \
    --engine=vcfeval \
    --engine-vcfeval-template {input.sdf} \
    {input.truth_vcf} \
    {input.query_vcf}
```

Here is a description of each parameter:

1. **threads** - The number of threads the process is allowed to use, 16 was used for all tests.
2. **params.out_prefix** - Output prefix
3. **input.reference** - Input reference FASTA, GRCh38 across all tests
4. **input.call_regions** - Input benchmark confidence regions
5. **params.sex** - Indicates if sample is male or female
6. **input.sdf** - Pre-processing file for the reference genome, which reduces run-times when the same reference is used repeatedly
7. **input.truth_vcf** - Input truth VCF file
8. **input.query_vcf** - Input query VCF file

##### 1.6.2 Aardvark

Aardvark was run using a static binary built for Linux with the same snakemake pipeline. The following command contains the exact paramterized options used to run Aardvark:

```
{input.binary} \
    compare \
    --threads {threads} \
    --reference {input.reference} \
    --truth-vcf {input.truth_vcf} \
    --query-vcf {input.query_vcf} \
    --regions {input.call_regions} \
    --min-variant-gap {input.min_variant_gap} \
    --output-dir {output.output_folder} \
    --output-debug {output.debug_folder}
```

Here is a description of each parameter:

1. **input.binary** - The input Aardvark binary file, which was the 0.7.4-static release available on the GitHub page.
2. **threads** - The number of threads the process is allowed to use, 16 for all tests.
3. **input.reference** - Input reference FASTA, GRCh38 across all tests
4. **input.truth_vcf** - Input truth VCF file
5. **input.query_vcf** - Input query VCF file
6. **input.call_regions** - Input benchmark confidence regions
7. **input.min_variant_gap** - Minimum gap between variants to group into a sub-problem (i.e. "window size"). For small variant only comparisons, we used the default of 50 bp which is sufficient for most use cases (see Section 2.7). When including SVs or STRs, we increased the window size to 1000 bp which improves the resulting metrics. This is discussed more in the main manuscript.
8. **output.output_folder** - Output folder for primary metrics and VCF file
9. **output.debug_folder** - Optional output folder for more detailed metrics

#### 2 Additional analyses

##### 2.1 Additional compute resource metrics

Figure 1 shows distributions of wall clock time, CPU time, and peak memory consumption for all runs of Aardvark. They are separated into runs with and without stratification, as stratification increased the compute requirements.

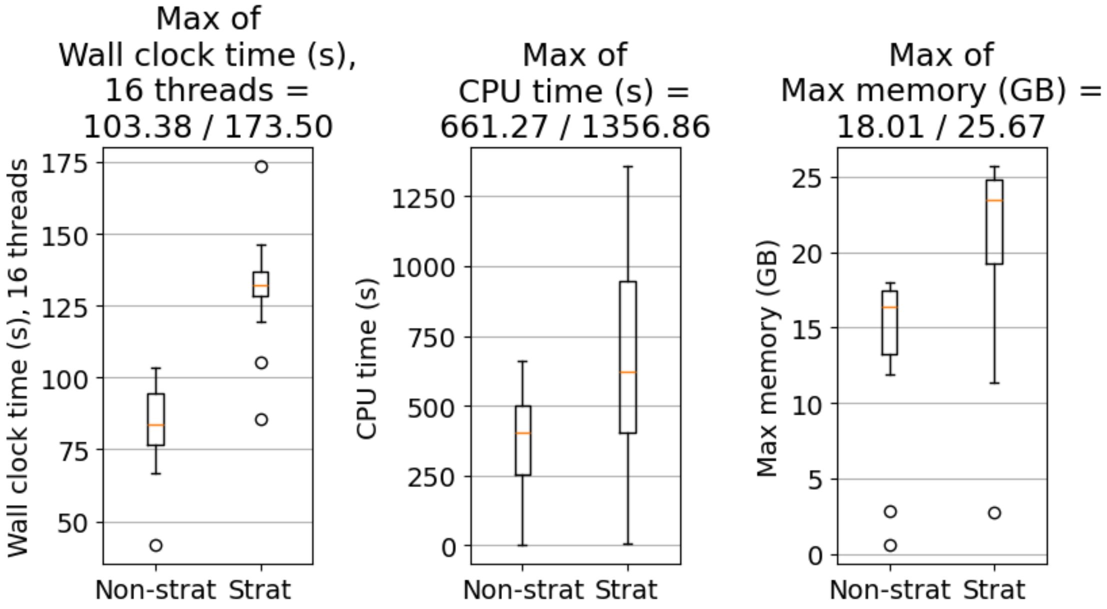

Figure 1: Compute metrics for Aardvark runs. These three figures show the distributions of wall clock time (16 threads), compute (CPU) time, and peak memory consumption across all runs of Aardvark. The data points are separated by whether stratification was run or not. Stratification increases compute and resource requirements across the board. Maximum values for each distribution are shown above the corresponding figures.

##### 2.2 Aardvark-GT has high concordance with hap.py

Aardvark's genotype scoring scheme has high concordance with hap.py (99.63-99.99%, see Figure 2), indicating that the vast majority of variants receive identical labels regardless of the input benchmark or technology used in the comparison. Concordance rates are highest for the GIAB benchmarks (>99.94%), which are more focused on low-complexity regions. In contrast, benchmarks that include higher complexity regions (specifically, CMRG and T2T) tend to have decreased concordance between Aardvark and hap.py.

The majority of discordant labels are caused by an intentional change in how genotype errors are counted in Aardvark relative to hap.py. Specifically, if there is a mismatch in the zygosity of a variant call (i.e., 0/1 in truth and 1/1 in query, or vice-versa), then hap.py will label both the truth and query variants as false. In contrast, Aardvark will only label the homozygous call as false, which both enables partial credit and is more consistent with Aardvark's other scoring approaches. This type of discordance occurs across all benchmarks and is the dominant form of discordance in most of our comparisons (see Figure 2).

Through manual inspection, we discovered another source of discordance around a representation of multi-allelic variants that is seemingly unsupported in hap.py and occurs in only the CMRG and T2T benchmarks. While these multi-allelic variants are almost always filtered from our discordance calculations, they often create a cascading effect causing nearby unfiltered variants to received a false positive/negative label from hap.py. This multi-allelic representation is supported in Aardvark, leading to increased discordance around multi-allelic sites with Aardvark assigning more true positive labels (see Supplemental Materials). We did

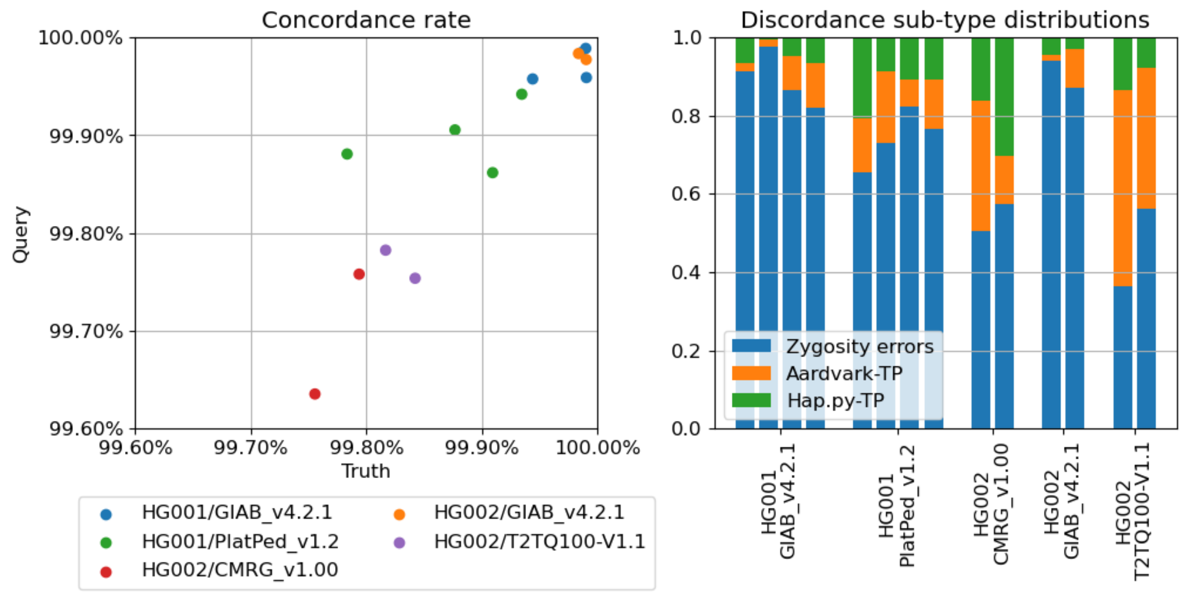

Figure 2: Concordance between Aardvark-`GT` and hap.py across different truth sets. Left: The overall concordance rate, indicating the fraction of variants in the truth (X-axis) and query (Y-axis) datasets with benchmark decisions ("BD" tag) that are the same in hap.py and Aardvark. Right: Distribution of discordant calls grouped by benchmark and type of discordance. "Genotype errors" are intentional reporting differences when the variant sequences match but are genotyped incorrectly (i.e., heterozygous v. homozygous). "Aardvark-TP" and "Hap.py-TP" are all other forms of discordance, indicating which tool labeled the discordant variant as a true positive.

not identify any other enriched categories of discordance. Critically, our manual inspections did not identify any clearly mis-labeled variants in the Aardvark results. Across all types of discordance, Aardvark tends to identify more true positives than hap.py. This culminates in slightly increased F1 scores across all comparisons for Aardvark's genotype score relative to hap.py (mean delta: +0.0016).

#### 2.3 Manual inspection details

We have created several miniaturized versions of interesting examples encountered while analyzing differences in hap.py and Aardvark. Some of these examples are pure mock examples, which usually include extremely simple reference sequences and basic variation to highlight a key difference between the two tools. Others are real examples (often labeled as "real_example_###"), which are a mixture of manually and automatically converted regions from our comparisons. The automatic conversions include notes on the sample, benchmark, dataset, and region that the example was copied from. For the most part, these miniature problems generated identical results in the real data and the miniaturized problem. However, we did find some examples that behaved differently in hap.py, which may be caused by some unknown subtle difference generated by our miniaturization methods.

The examples were all analyzed using a snakemake pipeline that runs hap.py, runs Aardvark, generates an MSA visualization of the Aardvark optimized haplotype sequences, and creates a summary README with all the metrics and additionally manually generated notes. All examples are available through this GitHub URL: `https://github.com/holtjma/mini_variant_benchmarks`.

The following are simple examples that may help with initial familiarization to the outputs:

1. simple_snv_het - A simple heterozygous SNV
2. simple_snv_hom - A simple homozygous SNV

##### 2.3.1 Examples of partial credit variants

The following are examples where one or more variants are partially correct at the BASEPAIR level, including many examples representing the common homopolymer calling error:

1. half_correct - A simple SNV / indel pair that do not exactly match up
2. half_correct_rev - The reverse example of the above
3. homopolymer_001 - Mock example with a homopolymer indel error
4. homopolymer_002 - Mock example with a homopolymer indel error
5. homopolymer_003 - Mock example with a homopolymer indel error
6. real_example_001 - Real example of partial credit
7. representation_mismatch_001 - Mock example with alternate representations and an error
8. representation_mismatch_002 - Mock example with alternate representations and an error
9. representation_mismatch_003 - Mock example with alternate representations and an error

##### 2.3.2 Examples of unsupported multi-allelics

The following examples are derived from real regions where hap.py labeled variants as "halfcalls" or otherwise seemingly ignored or skipped a multi-allelic variant. Interestingly, none of these examples actually replicated the "halfcall" labels that we saw in the full runs. We do not know the source of this difference.

1. real_example_006 - Unsupported "*" representation
2. real_example_007 - In our original dataset, this region did not resolve with hap.py. However, the mock version seemingly works, suggesting that perhaps our normalization has removed some representation that hap.py does not support. We are unclear of the discrepancy between the real and mock example.
3. real_example_008 - Multiple variants with "*" representation

##### 2.3.3 Examples with variant churn

As discussed in Section 3.4.1, we identified several examples of "variant churn", which we define as the presence of variants that nullify or counteract each other entirely or in part. We include some simple mock examples of variant churn in the first few examples below. Many of our manually inspected examples ("real_example_###" below) have some small amount of variant churn, indicating that they may be a major subtle source of discrepancy between hap.py and Aardvark.

The following are examples that include variant churn, including several real examples from our manual inspection:

1. null_variants_001 - Simple mock example
2. null_variants_002 - Simple mock example
3. null_variants_003 - Mock example with a true SNP and two null indels
4. real_example_003 - Example where two indels have a reduced representation
5. real_example_004 - Example where an indel and SNP combination have a reduced two indel representation
6. real_example_005 - Example where two indels have a reduced SNP representation
7. real_example_006 - Example where there are 8 total basepair changes in the truth set, but that reduces to 7 in sequence space, indicating the presence of complex variant churn

8. real_example_007 - High complexity region with variant churn in the truth set; 29 bp of changes expected from SNP and indels separately, but only 25 bp of sequence change when combined

9. real_example_008 - Complex example with 1 bp of variant churn in the truth set

10. real_example_009 - Complex example with 1 bp of variant churn in the truth set

11. real_example_012 - Complex example with 1 bp of variant churn in the truth set

12. real_example_013 - Complex example with 2 bp of variant churn in the truth set

#### 2.4 Comparison of all Aardvark scoring schemes

Figure 3 shows how the different types of scoring relate to each other relative to hap.py. Relative to hap.py, Aardvark genotype score tends to score higher due to increased ability to resolve complex variant representations and the ability to handle some types of multi-allelic variation that are unsupported in hap.py. Next, the haplotype scoring scheme evaluates each non-reference allele independently, effectively assigning equal weight to each alternate allele. The haplotype scoring scheme has slightly increased F1 scores relative to genotype scoring for both SR and LR query sets. The weighted haplotype scoring scheme additionally applies a weight to each alternate allele based on the length of the change, which enables each altered basepair to have equal weight in the scoring as with our basepair scoring scheme. In our query sets, the F1 scores are consistently lower for indels compared to SNVs, so implicitly giving more weight to indels leads to a drop in F1 score across all tests. If we focus on just the indel F1 scores, we actually see an increase in mean F1 scores for LR query sets (+0.0232) but a decrease for SR query sets (−0.0178, see Figure 3). This pattern is caused by our SR query sets excelling at shorter indels (≤15 bp) but struggling with longer indels (>15 bp), an observation that is supported by both hap.py and Aardvark-`GT` labeled outputs (see Section 2.5). These longer indels alter more basepairs and therefor have increased relative weight, leading to the relative drop in indel F1 scores for the SR query sets. We note that this trend persists when using SR and LR query sets from the Precision FDA Truth Challenge v2 [8] (see Section 2.6). Lastly, while both weighted haplotype and basepair scoring equalize the weight of each altered basepair, the basepair scoring scheme uniquely allows for partial credit when the variant sequences do not exactly match (see Table 7). Thus, basepair scores are expected to be strictly higher than the weighted haplotype scores, which is supported by each test in our analysis when all variants are considered jointly (mean delta: +0.0054).

#### 2.5 Stratification of indel F1 scores

In the main manuscript, we commented that the sequencing type has a major impact on the score differences between GT and BASEPAIR scoring schemes. We identified that the core reason for this is variability in indel detection depending on the length of the indel, which can be seen in the difference between HAP and WEIGHTED_HAP scoring schemes of Figure 3. In particular, short-read sequencing technologies seem to perform better with shorter indels, whereas long-read sequencing technologies seem to perform better with longer indels. Figure 4 shows this stratification using results from hap.py's genotype labels provided in the output VCFs. We see that the short-read technologies (Illumina and Element) tend to have higher F1 scores up to 20bp indels, but then drop rapidly in a bell-curve-like shape. In contrast, the long-read technologies (ONT and PacBio) tend to have a lower initial F1 score for smaller indels, but the curves are much flatter, leading to F1 scores that are higher than the short-read technologies for larger indels. While there are fewer of the larger indels, they impact more bases and therefor have a higher impact on the WEIGHTED_HAP and BASEPAIR scoring schemes. In contrast, the GT or HAP scoring schemes weight large indels the same as small indels, which reduces their impact on the scoring. While Figure 4 is derived purely from hap.py genotype labels, we notice similar trends when we use the Aardvark labels, indicating that this is unlikely an artifact from either tool. We suspect this trend is caused by challenges from alignment of short read fragments that may contain the evidence for these larger indels, which then cascade into challenges identifying these larger indels in the variant calling steps.

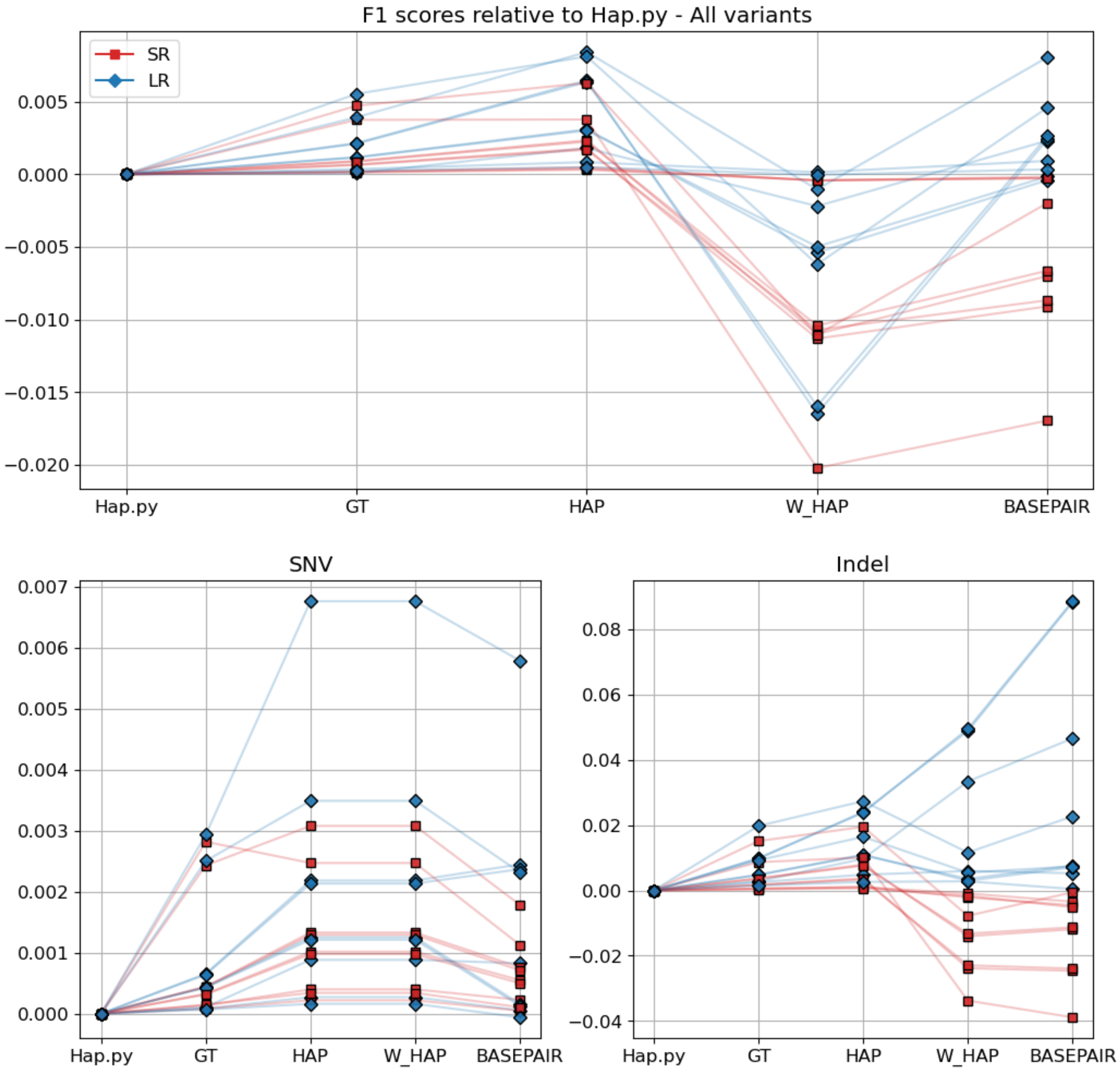

Figure 3: Aardvark F1 scores relative to hap.py. These figures shows the progressive difference in F1 scores relative to hap.py for all variants (top), SNVs (bottom left), and all indels (bottom right) when moving from a genotype-centric approach (hap.py and GT) to a sequence-centric approach (BASEPAIR). The marker shape and color indicates whether the query dataset is from short-read (SR) or long-read (LR) sequencing. Aardvark's genotype (GT) scoring is most similar to hap.py, assigning equal weight to each non-reference genotype call regardless of zygosity or length of the change. Aardvark's haplotype (HAP) score instead evaluates each non-reference allele independently, effectively doubling the weight of homozygous variant calls and allowing for partial credit if the genotype zygosities mismatch. Aardvark's weighted haplotype (W_HAP) scoring differs from the haplotype scoring scheme by weighting each allele by the length of the change, equalizing the weight of each altered basepair and implicitly assigning greater weight to alleles that impact more sequence (i.e., larger indels). Lastly, basepair (BASEPAIR) scoring evaluates full length haplotype sequences in a variant-agnostic approach, which allows for implicit partial credit when allelic sequences do not exactly match.

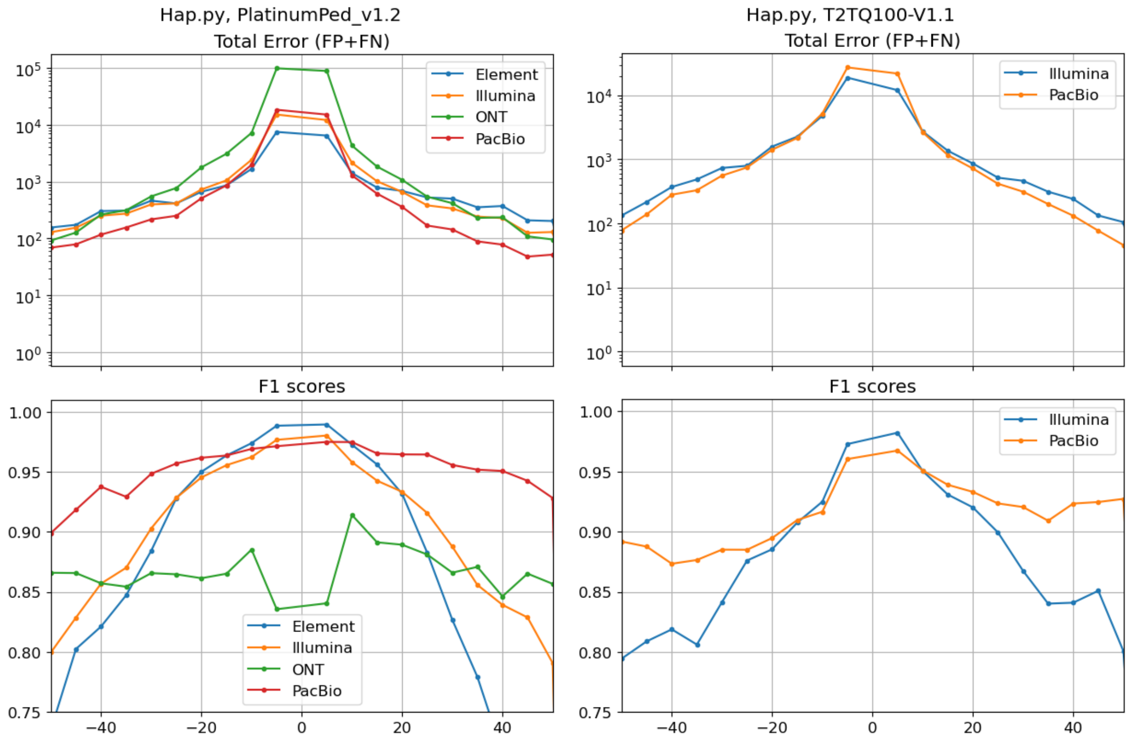

Figure 4: Indel stratification from hap.py. These figures show the indel stratifications grouped by indel sizes (grouped in 5 bp windows) for both HG001-PlatPed v1.2 (left) and HG002-T2T (right). Negative values indicate deletion variants, and positive values indicate insertion variants. The top figures show the total number of false positive (FP) and false negative (FN) labels in the hap.py output file. The bottom figures show the F1 scores for the corresponding indel size range. In general, the short-read technologies have a higher F1 score for shorter indels, but drop off quickly as the indels get larger. In contrast, the long-read technologies tend to have a lower initial F1 score for the shorter indels, but the drop is much slower as the indels get larger, leading to higher F1 scores for the larger indels.

#### 2.6 Precision FDA Truth Challenge v2 score trends

We repeated our analysis of the scoring schemes for variant calls that were submitted to the Precision FDA Truth Challenge v2 [8], comparing all HG002 benchmarks using Aardvark only. We note that some of the submissions failed to complete through Aardvark due to file formatting, and they are excluded from this analysis. All VCFs were downloaded following the instructions in [8], with the metadata pull from the GitHub repository[17].

Figure 5 shows the F1 scores relative to Aardvark-GT for the datasets that successfully completed the process, again stratified by the underlying sequence technology type (Illumina, ONT, PacBio, or "MULTI" for multiple technologies). In general, the trends observed in our main data persist here. Namely, the short-read entries have a drop in indel F1 score for the basepair scoring scheme, whereas long-read entries tend to increase. This is particularly true for the ONT submissions, showing consistent gains $\geq 0.05$. We also note that this trend first manifests when moving from haplotype to weighted haplotype scoring, indicating that the change is likely a direct result of weighting each modified base equally.

#### 2.7 Impact of window size on small variant benchmarking

Aardvark's window size is controllable with the `--min-variant-gap` parameter. Conceptually, this is the minimum gap in basepairs between two consecutive truth or query variants required to split into two separate subproblems. If the window size is too small, then the sub-problems may be too fractured, leading to increased false positives/negatives because matching variants have been placed in separate sub-problems. This is particularly true if the truth and query sets were generated with approaches that frequently represent variants differently (e.g., different technologies or variant callers). As the window size gets larger, the sub-problems get larger and the likelihood of over-fracturing the problems decreases, but the complexity of the problems increases which is expected to increase compute requirements. By default, Aardvark's window size is 50 bp. For context, the comparable window size parameter in hap.py is set to 30 bp by default.

Figure 6 shows the impact of increasing this parameter in Aardvark for small variants. In general, we see relatively little impact on overall F1 score as this parameter increases, potentially with some slight benefits up to 200 bp. Run-time (both wall clock and CPU time) does not change significantly until about 400 bp, where it begins to increase rapidly. Surprisingly, the max memory requirements decrease until 400 bp, reflecting that the bulk of the memory cost is likely caused by overhead from tracking sub-problems and not from the underlying algorithmic complexity. After 400 bp, it increases rapidly, which likely reflects the algorithm complexity from large sub-problems becoming the dominant driver of memory consumption. Overall for small variant benchmarking, these figures suggest that there may be a slight benefit to increasing the window size to 200 bp or so, without a major performance penalty. However, these tests were run using default settings (other than the window size), and do not reflect additional computations that users may frequently include (specifically, `--debug-folder` is not active). Additionally, when including larger SVs or STRs, we noticed higher F1 scores when using a window size of 1000 bp indicating that the types of variation present may also influence the optimal window size choice.

#### 2.8 Stratified difference in GT and BASEPAIR

When we stratify the results using the GIAB stratifications, we can hone in further on regions that benefit from a basepair scoring scheme. In particular, regions with tandem repeats or homopolymers are more prone to minor errors in sequence and/or variant calling. These regions benefit the most from basepair scoring, while those without these features do not (see Figure 7). We recommend developers who are looking to improve their tools assess the results from Aardvark with stratifications to better isolate problematic regions.

[17] `https://github.com/usnistgov/giab-pFDA-2nd-challenge/blob/cfcf6b12f1c28c8cbabfc5f6d671c18e670640e7/data-raw/anonymized_metadata_table.tsv`

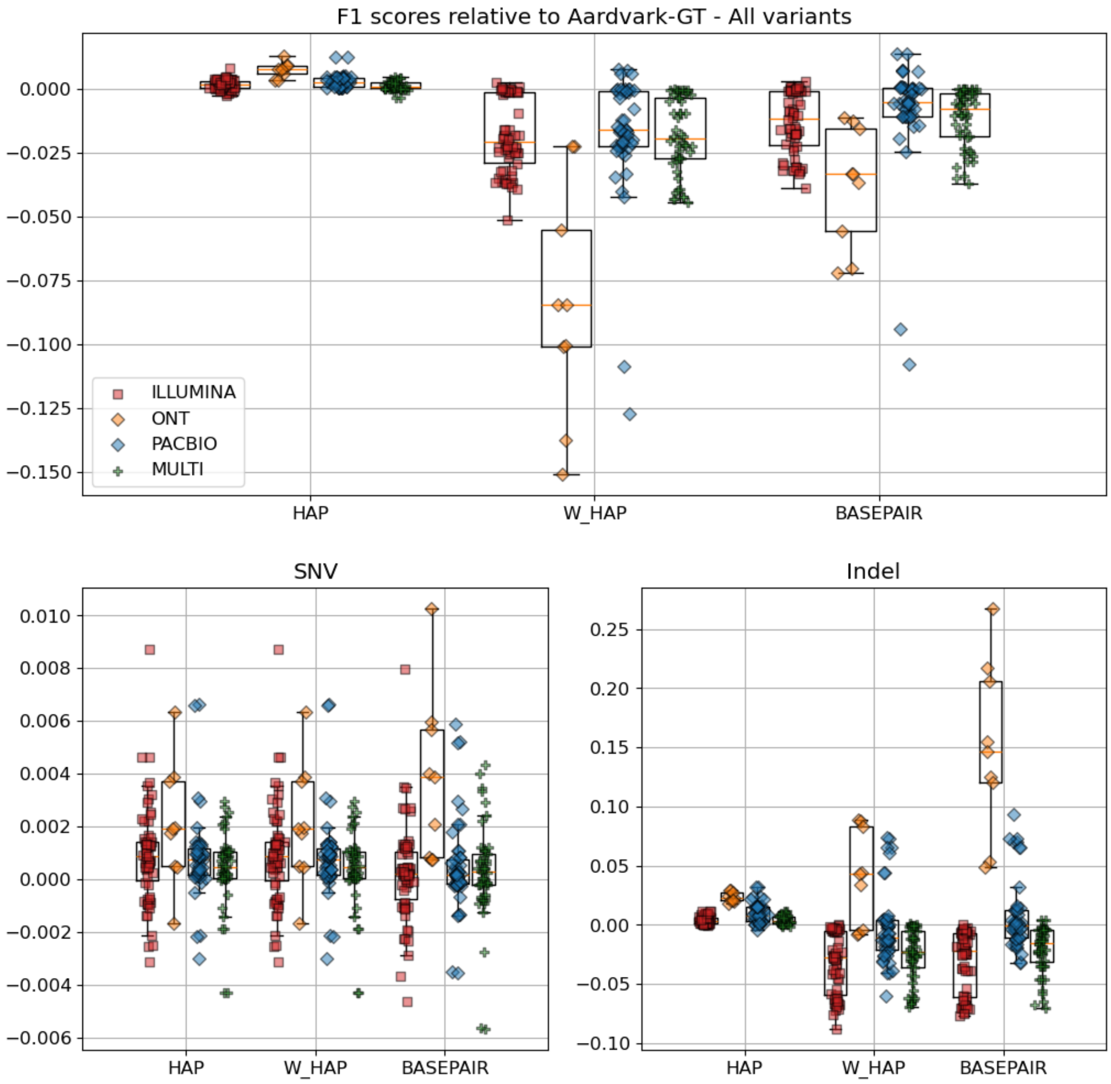

Figure 5: Aardvark F1 scores for Precision FDA Truth Challenge v2 HG002 submissions. The submissions were all run through Aardvark using an identical process to the data in the main document, comparing each entry to all three HG002 benchmarks. Data is plotted relative to Aardvark-GT's F1 score, which are usually slightly higher than those from hap.py. They are grouped by the types of sequencing used to generate the VCF file: Illumina (SR), ONT (LR), PacBio (LR), and "MULTI" which are generated from two or more of the other data types. These datasets support the same observation from the main document, where short-read variant calls tend to have a drop in indel F1 score for the basepair scoring. This drop is also reflected in the weighted haplotype (W_HAP) scoring scheme. In contrast, the long-read datasets tend to have higher or equal basepair F1 scores, with ONT-based variant calls showing the largest average gain relative to genotype scoring.

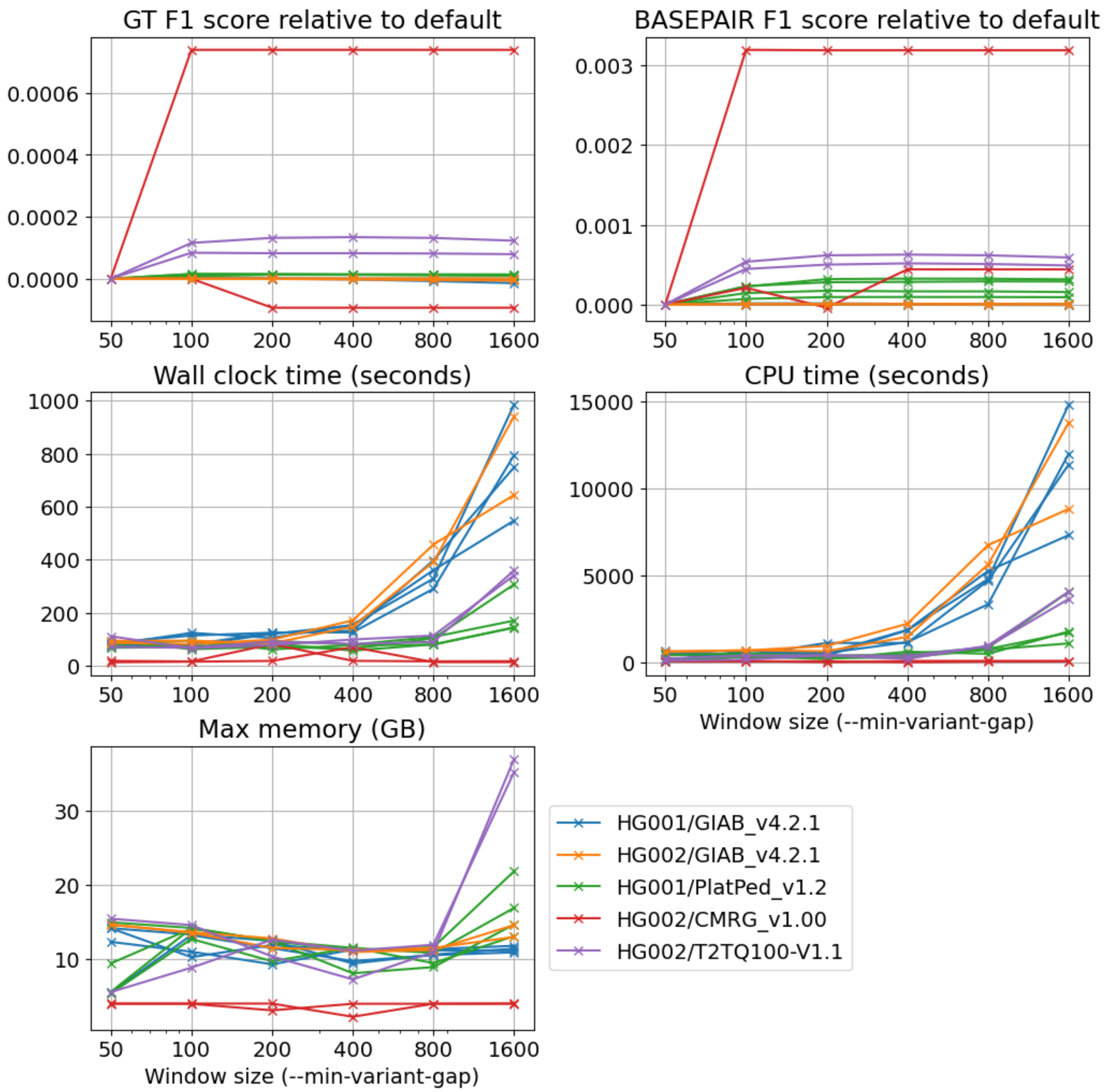

Figure 6: Impact of window size on Aardvark. These figures shows the impact of increasing the window size (`--min-variant-gap`) in Aardvark beyond the default of 50 bp. Overall, we see relatively little impact on F1 scores, with some slight benefits up to 200 bp or so. Additionally, compute times are relatively flat up to 200 bp, with a rapid increase beyond that. Max memory decreases until 400 bp, reflecting that the bulk of memory costs are likely sub-problem overhead up to that point, and then increasing rapidly as the underlying algorithmic complexity becomes the dominant driver of memory usage.

### 3 Methods details

#### 3.1 Summary of genotype scoring critiques

Table 4 summarizes a collection of critiques that can be made of genotype scoring, along with the way basepair scoring alleviates or reduces the issue. These are summarized in the discussion of the main manuscript, but the table is useful for highlighting particular critiques and clustering them with similar issues.

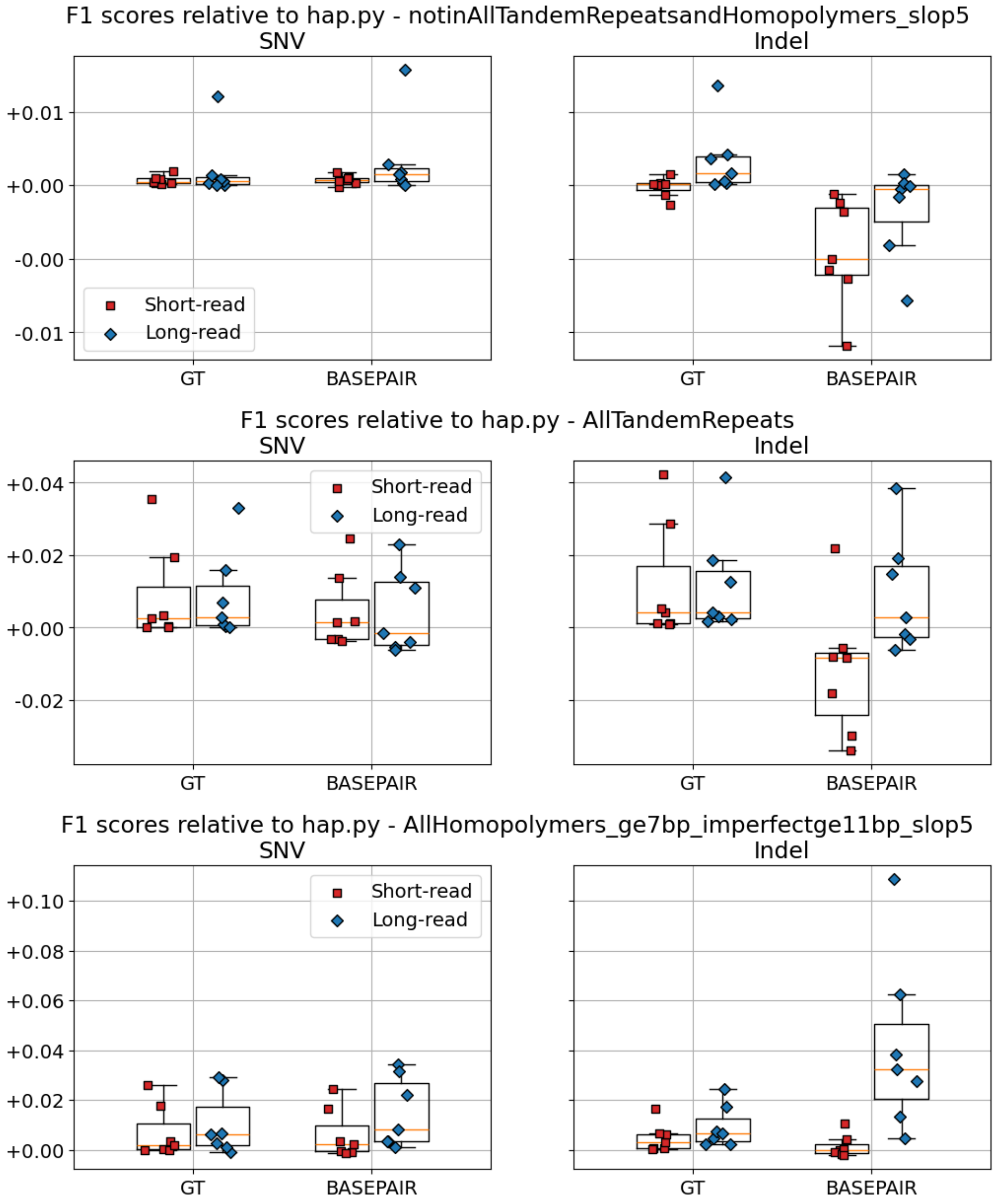

Figure 7: F1 scores in stratification regions. These figures show the F1 scores relative to hap.py for SNVs (left) and indels (right) across three stratification regions. The first region excludes tandem repeat and homopolymer regions, the second includes only tandem repeats, and the last includes homopolymers. We observe that outside of tandem repeat and homopolymer regions, we see a decrease in the basepair scoring relative to hap.py. In contrast, the basepair score tends to be higher inside tandem repeat and homopolymer regions, especially for the long-read technologies.

<table>
<tr><th>Critiques of genotype scoring</th><th>Differences in basepair scoring</th></tr>
<tr><td>Variant representation for equivalent haplotypes can influence the total weight</td><td rowspan="3">Variant representation is masked and only the haplotype sequences represented by the variants are compared</td></tr>
<tr><td>Variant representation may be influenced by technology, aligner, or variant caller, generating a bias for or against some combinations</td></tr>
<tr><td>Specialized callers (e.g., tandem repeat callers) cannot be compared to a small variant benchmark because the variant representations are too different</td></tr>
<tr><td>Variants of different lengths have the same weight, despite one of them impacting more sequence (e.g., a 1-bp indel carries the same weight as a 50-bp indel)</td><td>Variant calls carry weight proportional to the number of bases they modify, so longer variants will carry more weight</td></tr>
<tr><td>Homozygous variants have equal weight as heterozygous variants despite impacting twice the genomic sequence</td><td rowspan="2">Homozygous calls implicitly carry twice the weight of a heterozygous call since they impact both haplotype sequences, and zygosity errors are given partial credit (e.g., 50%)</td></tr>
<tr><td>Zygosity errors (e.g., 0/1 instead of 1/1) count the same as missing the variant entirely (e.g., 0/0 instead of 1/1)</td></tr>
<tr><td>Off-by-one errors in the alleles will get no credit despite getting the variant mostly correct</td><td rowspan="2">Each correct basepair is counted as a true positive and all others as false positive/negative, enabling partial credit when alleles are not exact matches</td></tr>
<tr><td>No-calls are treated the same as a partially-correct allelic sequence</td></tr>
</table>

Table 4: A list of critiques of genotype scoring, which are grouped into categories. Beside each grouping is the key differences with basepair scoring that overcome these critiques in part or entirely.

#### 3.2 Variant-based additional scoring schemes

In the main document, we focused on the genotype and basepair scoring schemes as these are the primary scores from Aardvark. However, Aardvark includes two additional variant-base scoring schemes that are derived from the same calculation used to generate the genotype results. These alternative schemes are helpful in understanding the progression from a genotype-centric scoring scheme to sequence-centric scoring schemes. We are also exploring other scoring schemes that may be more appropriate for tandem repeats or structural variants, but these are not yet implemented.

##### 3.2.1 Genotype scoring scheme

For the genotype scoring scheme, Aardvark starts by using the phase orientations from sequence optimization, and independently evaluates each pair of truth and query alternate alleles. For each pair of alternate alleles, Aardvark searches for the maximum number of alternate alleles that can be incorporated into the reference sequence such that the output truth and query haplotype sequences exactly match. Critically, the allele representations do not have to match, so long as the final haplotype sequences are basepair-identical. Any alleles that are incorporated are marked as true positives, and any that are not are either false negatives or false positives depending on the source. For homozygous genotypes, the genotype score further requires that both alleles are marked as true positives for the entire genotype to also get marked as a true positive. This definition is consistent with previous approaches [3, 6].

Similar to the sequence optimization problem, Aardvark models this process as a decision tree where at depth $A$, the tool is deciding if allele $A$ is true or false. As it traverses the tree, it extends the truth and query haplotypes with any true alleles, removing the node from future consideration if any mismatches are identified (i.e., at least one of the included true alleles should be false). When the search reaches depth $A$, each allele up to depth $A$ has been labeled as either true or false, with all true alleles forming truth and query haplotype sequences that contain no mismatches. This problem can have multiple solutions (e.g., all FP/FN is always a valid solution), so Aardvark searches for the solution with the most true alleles in the output (i.e., maximize the number of true positive labels). Fortunately, clever optimizations introduced by `vcfeval` allow for "checkpoints" in the search that significantly reduce the search space in practice [3].

There are two subtleties worth mentioning about the Aardvark implementation of the genotype score. First, any multi-allelic heterozygous sites are split into two distinct heterozygous genotypes for all analyses. Second, if the difference between truth and query is limited to the zygosity of the genotype call (i.e., heterozygous v. homozygous), then the heterozygous call is labeled as "true" and the homozygous call is labeled "false". For example, if the truth set has a heterozygous call while the query set has homozygous, then the truth variant is labeled as a true positive since it was correctly detected in the query, but the query variant is labeled as a false positive since an extra allele is present. Both of these subtleties are distinctions from the hap.py genotype scoring [6], and are sources of technical discrepancy when comparing the two tools. We include examples of each subtlety in Table 5.

##### 3.2.2 Haplotype scoring schemes

Aardvark reports two additional scores that are derived from the same calculations used to generate the genotype scores, and serve as intermediate scores on the way to a full basepair scoring scheme. First, the haplotype (`HAP`) score evaluates each alternate allele in the truth and query sets independently. For heterozygous calls, the scoring is identical to genotype scoring as only a single alternate allele is present. For homozygous calls, each alternate allele has a weight of 1 and is evaluated independently. This change equalizes the impact of each alternate allele such that one homozygous variant has the same weight as two heterozygous variants, and it also allows for partial credit when only the zygosity errors occur in the genotypes (i.e., 0/1 v. 1/1). Conceptually, this scoring scheme moves from a recall/precision denominator that is the number of genotypes to one that is the number of alternate alleles.

Second, the weighted haplotype (`WEIGHTED_HAP`) scoring scheme is calculated the same as the haplotype score, except each allele is additionally weighted by the edit distance between the reference and alternate sequences. For SNVs, this has no impact on scoring since the edit distance by definition is 1. However, indel variants receive a weight based on how many bases they insert or delete, enabling a single 2-bp insertion to have the same weight as two separate 1-bp insertions. This further moves the recall/precision denominator

| Set | REF | Truth | Query | GT | HAP | W_HAP |
|---|---|---|---|---|---|---|
| A | A | A/C | A/C | TP=1 | TP=1 | TP=1 |
| | A | A/C | C/C | Truth.TP=1, FP=1 | TP=1, FP=1 | TP=1, FP=1 |
| | A | C/C | A/C | FN=1, Query.TP=1 | TP=1, FN=1 | TP=1, FN=1 |
| | A | C/C | C/C | TP=1 | TP=2 | TP=2 |
| B | A | A/ACC | A/ACC | TP=1 | TP=1 | TP=2 |
| | A | A/ACC | ACC/ACC | Truth.TP=1, FP=1 | TP=1, FP=1 | TP=2, FP=2 |
| | A | ACC/ACC | A/ACC | FN=1, Query.TP=1 | TP=1, FN=1 | TP=2, FN=2 |
| | A | ACC/ACC | ACC/ACC | TP=1 | TP=2 | TP=4 |
| C | A | ACC/ACCC | A/A | FN=2 | FN=2 | FN=5 |
| | A | ACC/ACCC | A/ACC | TP=1, FN=1 | TP=1, FN=1 | TP=2, FN=3 |
| | A | ACC/ACCC | A/ACCC | TP=1, FN=1 | TP=1, FN=1 | TP=3, FN=2 |
| | A | ACC/ACCC | ACC/ACCC | TP=2 | TP=2 | TP=5 |

Table 5: Examples of variant-based Aardvark scoring schemes. The examples in this table include the reference sequence (REF), the Truth genotype call, the Query genotype call, and the generated number of true positives (TP), false negatives (FN), and false positives (FP) for Aardvark's genotype (`GT`), haplotype (`HAP`), and weighted haplotype (`W_HAP`) scoring schemes. Set A are all simple SNVs, emphasizing the scoring differences in the `GT` and `HAP` scores when genotypes are different and all alleles have a weight of 1. Set B changes to an 2-bp insertion variant, highlighting that all `W_HAP` scores now have twice the total weight of the `HAP` scoring. Set C highlights how multi-allelic genotypes are treated as two distinct genotypes across all scoring schemes, while also showing how alternate alleles of different lengths are scored in the `W_HAP` scheme. FN and FP are always unique to truth and query, respectively. TP may be different between truth and query (impacting recall and precision) and is annotated as "Truth.TP" and "Query.TP" where relevant. If TP, FN, or FP is unlisted, it is =0.

from alternates haplotypes to number of altered basepairs. We provide several examples of differences in the genotype, haplotype, and weighted haplotype scoring in Table 5.

The main benefits of the haplotype and weighted haplotype scoring schemes are reduced biases implicitly caused by variant representation. In particular, indel variants can often be represented in multiple ways, which may significantly alter both the number of variant entries in a VCF file and their zygosities. Table 6 shows examples where the representation can skew the relative weights of altered basepairs, which can lead to significantly different results even when the underlying haplotypes are sequence-identical.

#### 3.3 Basepair v. weighted haplotype scoring schemes

While the weighted haplotype scoring scheme resolves many representation bias issues, it does not allow for partial credit when alleles nearly match. This can lead to scenarios where the resulting metrics may still depend on the representations chosen by truth and query. The basepair scoring alleviates this last issue, and we highlight differences between weighted haplotype and basepair in Table 7.

#### 3.4 Basepair variant-type scoring

The basepair scoring scheme is intended to be a comprehensive score that factors in all provided variants, and removes their representations from consideration. However, there is often a desire to distinguish the accuracy of variants by their type (i.e., SNV or indel), which is not possible when those representations have been removed. To get around this limitation, Aardvark will actually perform multiple passes of the basepair scoring, but with different haplotype sequences formed from subsets of the full variation. In particular, it will create a haplotype sequence using only one variant type and then compare that sequence to the full sequence from the other source. For example, it may compare a query sequence with only SNVs against a truth sequence with all variants included. This specific comparison will generate many false negatives since only a subset of the query variation is present, but the algorithm knows that any identified true positive or false positive bases must come from query SNVs (and not other variant types). The process is repeated for each variant type in the query set. Then, a reciprocal process is used where the query variants are all

| Set | Variants | Altered bp | GT | HAP | W_HAP | BASEPAIR |
|---|---|---|---|---|---|---|
| A | One Het. SNV | 1 | 1 (0.0) | 1 (0.0) | 1 (0.0) | 1 (0.0) |
| B | Two Het. 1-bp indels | 2 | 2 (0.5) | 2 (0.5) | 2 (0.5) | 2 (0.5) |
| | One Hom. 1-bp indel | 2 | 1 (0.0) | 2 (0.5) | 2 (0.5) | 2 (0.5) |
| C | One Het. 50-bp indel | 50 | 1 (0.0) | 1 (0.0) | 50 (0.0) | 50 (0.98) |
| | Two Het. 25-bp indels | 50 | 2 (0.5) | 2 (0.5) | 50 (0.5) | 50 (0.98) |
| | Ten Het. 5-bp indels | 50 | 10 (0.9) | 10 (0.9) | 50 (0.9) | 50 (0.98) |
| | Fifty Het. 1-bp indels | 50 | 50 (0.98) | 50 (0.98) | 50 (0.98) | 50 (0.98) |
| D | One Hom. 50-bp indel | 100 | 1 (0.0) | 2 (0.5) | 100 (0.5) | 100 (0.99) |
| | Two Hom. 25-bp indels | 100 | 2 (0.5) | 4 (0.75) | 100 (0.75) | 100 (0.99) |
| | Ten Hom. 5-bp indels | 100 | 10 (0.9) | 20 (0.95) | 100 (0.95) | 100 (0.99) |
| | Fifty Hom. 1-bp indels | 100 | 50 (0.98) | 100 (0.99) | 100 (0.99) | 100 (0.99) |

Table 6: Detailed examples of representation biases. This table is a more detailed version of the table in the main document that includes the additional scoring schemes. This table shows multiple simplified examples of variant combinations that are labeled with a count, zygosity (heterozygous or homozygous), and variant type. The variant combinations are grouped into sets based on the number of altered basepairs (changed, inserted, or deleted), highlighting situations that could be sequence-identical. For each of the scoring schemes, the total weight of the variants under that scoring scheme is shown along with recall metric if a 1-bp false negative error were present in the query set. Set A has a basic heterozygous SNV, where all scoring schemes have identical weight. Set B shows two scenarios that equal 2 basepairs of inserted sequence, emphasizing the differences in weighting between the genotype (`GT`) and haplotype (`HAP`) scoring schemes. Sets C and D show four scenarios each that equal a 50 bp heterozygous or homozygous indel, further emphasizing major differences in weighting between the approaches. For the `GT` scoring scheme, the total weight is always equal to the number of variant entries from the VCF file. In contrast, we see that the weighted haplotype (`W_HAP`) and basepair (`BASEPAIR`) scoring schemes both have a total weight equal to the number of altered basepairs. Note that the `GT`, `HAP`, and `W_HAP` recall values are heavily influenced by the variant representation. In contrast, all recall values for the `BASEPAIR` scoring scheme are identical within each set, indicating that the representation has no impact on the final scoring. Overall, the `BASEPAIR` weighting and scoring does not change with the provided variant representations, removing many sources of bias that are present in the other scoring approaches.

| Set | REF | Truth | Query | W_HAP | BASEPAIR |
|---|---|---|---|---|---|
| A | A | A/C | A/C | TP=1 | TP=2 |
| | A | A/C | A/A | FN=1 | FN=2 |
| | A | A/A | A/C | FP=1 | FP=2 |
| | A | A/C | A/G | FN=1, FP=1 | TP=1, FN=1, FP=1 |
| | A | A/C | A/AC | FN=1, FP=1 | TP=1, FN=1, FP=1 |
| B | A | A/ACCC | A/ACCC | TP=3 | TP=6 |
| | A | A/ACCC | A/ACC | FN=3, FP=2 | TP=4, FN=2 |
| | A | A/ACCC | A/ACCCC | FN=3, FP=4 | TP=6, FP=2 |
| C | A | C/C | C/AC | TP=1, FN=1, FP=1 | TP=3, FN=1, FP=1 |
| | A | ACCC/ACCC | ACC/ACCCC | FN=6, FP=6 | TP=10, FN=2, FP=2 |

Table 7: Examples of differences between Aardvark's weighted haplotype (`W_HAP`) and basepair (`BASEPAIR`) scoring schemes. The examples in this table include the reference sequence (REF), the truth sequence, the query sequence, and the generated number of true positives (TP), false negatives (FN), and false positives (FP) for both `W_HAP` and `BASEPAIR` scoring schemes. To avoid floating-point numbers from "half-correct" base changes, `BASEPAIR` values are twice the number of altered basepairs. Set A are all heterozygous variants with a single base change, including two simple examples where the "half-correct" changes are relevant in basepair scoring (any odd number is a "half-correct" situation). Set B are all heterozygous insertions, showing major scoring differences when partial credit is applied in homopolymer indel regions. Set C includes a mix of heterozygous and homozygous genotype calls on alleles that reflect common errors in homopolymer variant calling. FN and FP are always unique to truth and query, respectively, but TP is shared in these particular examples. If TP, FN, or FP is unlisted, it is =0.

incorporated and subsets of the truth variants are used to construct the haplotypes, which allows the method to measure false negatives as well. The core limitation of this approach is that the full context has been reduced, which can lead to some unexpected outcomes, a phenomenon we call "variant churn".

##### 3.4.1 Variant churn

In our manual inspections, we identified many situations where variants would nullify or counteract each other when placed on the same haplotype. For example, a 1-bp insertion followed by a 1-bp deletion of the same base results in no change to the full sequence. These basepairs effectively disappear from the joint scoring without affecting the number of true/false bases identified. However, when considered separately, each of these manifest as false positives/negatives because they cannot be found in the comparator haplotypes and they are no longer getting removed by some other variant. This tends to lead to lower scores for the variant-type basepair metrics relative to the combined basepair metrics. For example, if you have a 1-bp insertion that is nullified by a 1-bp deletion, then nothing is contributed to the combined basepair scoring, but the variant-level metrics will show 2-bp worth of false positives/negatives.

While this may seem like a limitation, it actually provides the basis for a "variant churn" metric, which measures the fraction of bases in a VCF file that are unnecessary to describe the haplotypes compared in Aardvark. Critically, if all bases that are part of the variant churn are removed, the resulting haplotype sequences are still identical. With basepair scoring, we can estimate the amount of variant churn present within both the truth and query sets using the summary metrics file. We extract the total number of altered basepairs from the combined scoring, which represents the number of variant changes *after* null variants have canceled each other. We then sum the total number of altered basepairs from each separate variant category, which represents the total number of variant changes *before* null variants have canceled each other. Subtracting these numbers gives the total number of basepairs that are unnecessary to represent the sequence, and this value is the variant churn metric for a specific comparison. This value is calculated separately for both the truth and query sets, but we note that the exact number will depend on the truth set, query set, and region selection, which is why we refer to it as an estimate.

Figure 8 shows the measured churn as a fraction of the total input alternate basepairs for each run in our analyses. The runs are grouped by test and query sets, with each Aardvark comparison represented once in each sub-figure (once for truth, once for query). We note that most query sets have a relatively low churn fraction, < 0.5%, and that variation within the query points seems to be driven by the comparator truth set. In contrast, the truth sets seem to each exhibit their own variant churn value that is relatively consistent regardless of the chosen query set. For example, both GIAB truth sets have a consistently low variant churn fractions, < 0.2%, while the higher complexity benchmarks of CMRG and T2T have a consistently higher variant churn fractions, $1.0 - 1.4\%$. We hypothesize that this increase is driven by the inclusion of higher complexity regions, where variant churn is more likely to get generated by the pipelines and variant callers that were used to derive the truth sets. It is also possible that some amount of churn is expected due to variant representations that are biologically derived (as opposed to simplest representation), and perhaps that type of churn is enriched in the higher complexity benchmarks. For example, two adjacent homopolymers may be more likely to have independent indel variation, which by chance has a simpler SNV representation (e.g., ...AAA**A**CCC... to ...AAA**C**CCC...) that would manifest as variant churn in this analysis. It is unclear the degree to which each of these are occurring, as tracking the origin of the churn to specific variants is a non-trivial problem. Ideally, future work could detect and correct any null variation derived from technical artifacts in new versions of these benchmarks.

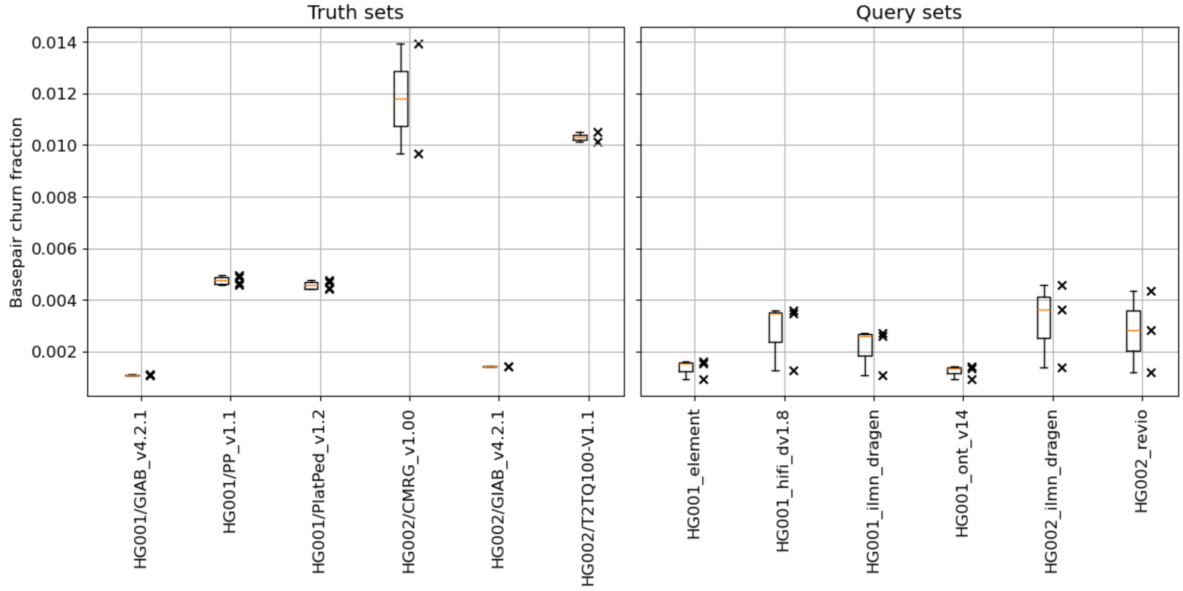

Figure 8: Variant churn fractions. These figures shown the calculated churn fraction for the test sets (left) and query sets (right). The churn fraction is defined as the fraction of bases that could be removed from the variant representations to achieve the same haplotype sequences. In these examples, we see that variant churn seems to vary by truth set significantly. Low complexity benchmarks (GIAB) tend to have low variant churn (< 0.2%) while higher complex benchmarks (CMRG and T2T) have a much higher churn fraction ($1.0 - 1.4\%$). Query sets do not show much difference in churn fraction, and variance within a query set seems to be driven by the comparator test set.

#### 3.5 Platinum Pedigree tandem repeat analysis

The Platinum Pedigree includes small variant and tandem repeat benchmarks, each of which has its own VCF file and benchmark regions (BED) file. For the experiment in our main document, we needed to restrict these benchmark files down to the regions with high confidence in both benchmarks that have equivalent sequence-level representations. First, we intersected the two sets of benchmark regions and kept only those for which the entire tandem repeat region shared, leaving us with ≈494k regions:

```
bedtools intersect \
```

```
    -a ${STR_REGIONS} \
    -b ${SMALL_REGIONS} \
    -wa -u -f 1.0 \
    | bedtools sort \
    > ${SHARED_REGIONS_BED}$
```

Second, we used an auxiliary function of Aardvark to identify any regions which were not sequence level identical in the small variants and tandem repeat variants. Aardvark includes a `merge` routine which is most useful for identifying regions that are sequence-identical in different VCF files and collapsing them into a single representation. Part of the output is a failed regions file (BED) that indicates any regions which are not sequence equivalent. For simplicity, we opted to remove any regions that were not 100% sequence identical on both haplotypes between these representations. We subtracted any regions that overlapped this failure file:

```
# perform the merge
aardvark merge \
    -r ${REFERENCE} \
    -i ${PLATPED_SMALL_VCF} \
    -i ${PLATPED_STR_VCF} \
    -b ${SHARED_REGIONS_BED} \
    -o ${MERGE_RESULT} \
    --min-variant-gap 1000 \
    --merge-strategy exact \
    --threads 16

# subtract the failed regions
bedtools subtract \
    -a ${SHARED_REGIONS_BED} \
    -b ${MERGE_RESULT}/failed_regions.bed.gz \
    -A \
    | bedtools sort \
    > ${IDENTICAL_REGIONS_BED}
```

The resulting `IDENTICAL_REGIONS_BED` was used to limit our comparison, containing ≈413k sequence-identical confidence regions. Both of our query sets (DeepVariant and TRGT callers) demonstrated identical basepair F1 scores on each of the Platinum Pedigree benchmarks, verifying that we successfully reduced the small variant and tandem repeat benchmarks to the regions which were sequence identical.

#### 3.6 Measuring compute requirements

All benchmarks where we collected compute requirements were run through a snakemake pipeline to establish systematic execution of the commands used in our standard workflow. These pipelines have an option to write compute usage to a "benchmark" file, which was used for both Aardvark and hap.py measurements. While this approach works without issue on standard command line tools like Aardvark, it does not always accurately measure CPU time or memory usage if a Docker or Singularity image is used as with hap.py. Thus, we were able to obtain wall clock time, CPU time, and memory usage for Aardvark (see Section 2.1), but only wall clock time for hap.py.

#### 3.7 Calculating concordance

Both Aardvark and hap.py output truth and query variants with benchmark decision labels ("BD" tag) indicating if the variant is classified as a true positive (TP), false positive (FP), or false negative (FN). However, the two tools have different approaches for normalizing variants and processing multi-allelic sites. For example, hap.py will often split variants into smaller components (e.g., a multi-nucleotide variant may become multiple SNVs), whereas Aardvark will not split the alleles. Additionally, Aardvark will split non-reference heterozygous sites into two distinct heterozygous calls (e.g., 1/2 becomes two 0/1 genotypes). Lastly, the normalizations methods for each tool are different, and often generate slightly different results for multi-allelic sites.

To identify these problematic sites, we merged the output VCF files from both hap.py and Aardvark, and then excluding any variants where the genotypes or allelic sequences did not match. This removed the variants that are normalized differently or that were split into multiple genotypes. The remaining variants were included in our concordance analysis, and we simply compared the benchmark decision labels to determine if they were concordant (matching labels) or discordant.

#### 3.8 Experimental joint variant calling pipeline

There are few variant callers that will produce an integrated VCF file containing small variants and larger forms of variation (tandem repeats or structural variants). Additionally, most of these accept an assembly as input, limiting the reported variants to only regions that are successfully assembled. Instead of using one of these assembly approaches, we used an experimental joint variant calling pipeline with the following steps:

1. Generate a diploid *de novo* assembly from long-read data - We used an HG002 HiFi sequence dataset as input. The assembly was generated with hifiasm [2] for the assembly.

2. Align the assembly to GRCh38 - We used minimap2 for this alignment.

3. Align the reads to the assembly - We used minimap2 to then align the original reads to the diploid *de novo* assembly.

4. Liftover the mapped reads back to GRCh38 - We used portello[18] to perform this liftover. Portello is an experimental software that performs liftover from assembly-mapped reads to a standard reference genome. This tends to create more consistent alignments around sequencing errors, improving downstream variant calling.

5. Jointly call small and structural variants - We used longcallD[19] to jointly call phased variants from the alignment. LongcallD is in pre-release state, but is one of the few variant callers that will produce a joint VCF. This final VCF is included in our data bundle.

[18] https://github.com/PacificBiosciences/portello
[19] https://github.com/yangao07/longcallD

### 4 Notes on benchmarking

#### 4.1 Tandem repeat catalogs

When benchmarking, it is essential to use balanced tandem repeat truth sets in which non-homozygous reference repeats are adequately represented. If all repeats are homozygous reference, even a single basepair error can drive precision to zero for both genotype and basepair scoring. This scenario is not merely theoretical, as it is common for genome-wide repeat catalogs to contain repeats that are homozygous reference across the majority of analyzed samples.

Historically, tandem repeat studies have reported the accuracy of alleles listed in the corresponding VCFs, irrespective of how closely their sequences match the reference genome. For example, if the reference allele in a tandem repeat region is 100 bp long and the truth set has the reference allele for a sample, then a 98 bp query allele would have a score of 0.98 (98 / 100). In contrast, the basepair precision for this same allele would show two false positive deletion bases, resulting in a precision score of 0.0 for the region. For tandem repeat callers that are reference-agnostic, this historical approach can be appropriate as the alleles are constructed without knowledge of the reference allele, with all final metrics based on the differences (e.g., edit distance) and normalized against the lengths of the compared alleles. However, if the reference alleles are known to the tandem repeat caller, then the resulting scores may be inflated due to prior knowledge and biases towards a reference allele. Thus, the exact repeat catalog chosen can influence the resulting metrics of this approach. As with genotype scoring, we consider this historical score to be complementary to Aardvark's basepair scoring, and using both may help identify areas to improve tandem repeat callers.
